# A temperature-tolerant CRISPR base editor mediates highly efficient and precise gene inactivation *in vivo*

**DOI:** 10.1101/2022.12.13.520203

**Authors:** Roman M. Doll, Michael Boutros, Fillip Port

**Affiliations:** German Cancer Research Center (DKFZ), Division of Signaling and Functional Genomics and Heidelberg University, Heidelberg, Germany; Molecular Biosciences/Cancer Biology Program, Heidelberg University and German Cancer Research Center (DKFZ), Heidelberg, Germany

**Author notes:** MRC Molecular Haematology Unit, MRC Weatherall Institute of Molecular Medicine, Radcliffe Department of Medicine, University of Oxford, Oxford, UK.

**Keywords:** Base editing, CRISPR, genome engineering, gene knock-out, Drosophila

## Abstract

CRISPR nucleases generate a broad spectrum of mutations that includes undesired editing outcomes which attenuate phenotypes and complicate experimental analysis and interpretation. Here, we develop an optimised cytosine base editing system for gene inactivation in *Drosophila* through predictable C-to-T editing and identify temperature as a crucial parameter for base editing efficiency. We find that activity of an evolved version of the most widely used APOBEC1 deaminase is attenuated within the temperature range commonly used for culturing *Drosophila* (18-29°C) and many other ectothermic species. In contrast, an evolved CDA1 domain functions with remarkable efficiency within the same temperature range. Furthermore, we show that formation of undesired indel mutations and C-to-G/A edits is exceptionally rare in *Drosophila* compared to other species. The predictable editing outcome, very high efficiency and minimal byproduct formation of this system allows for near homogeneous biallelic gene inactivation *in vivo* in a ubiquitous or conditional manner. This work significantly improves our ability to create precise loss-of-function alleles in *Drosophila* and provides key design parameters for developing highly efficient base editing systems in other ectothermic species.

## Introduction

The inactivation of genes through the induction of loss-of-function mutations is a central tool to establish causal genotype-phenotype relationships. Over the last decade, CRISPR-Cas systems have emerged as the preferred system for the installation of targeted mutations, mainly due to their high efficiency and ease of use. CRISPR nucleases, such as Cas9 or Cas12a, can be programmed by guide RNAs (gRNAs) to create DNA double-strand breaks (DSBs) at defined genomic loci, which are resolved by error-prone endogenous DNA repair pathways, thereby creating mutations at the target site^1–3^. A pervasive shortcoming of this strategy is the uncontrollable nature of the induced DNA alterations, which are typically dominated by short insertions and deletions (indels)^4^, but also include large indels and genomic rearrangements at significant frequency^5,6^. Typically, not all of the induced mutations have the desired effect. For example, small in-frame indels are often functionally silent when induced in the coding sequence of a gene^4,7,8^ and larger deletions and rearrangements can cause false-positive phenotypes by affecting genes other than the target. Therefore, methods that can efficiently generate precise mutations would offer advantages for gene inactivation experiments.

CRISPR base editors are fusion proteins of a Cas9 nickase linked to a deaminase domain and can be used to install defined point mutations in the genome^9–11^. Cytosine base editors (CBEs) create C-to-T edits through i) R-loop formation, ii) deamination of cytosine residues in the single-stranded, non-target DNA strand by a cytidine deaminase to convert cytosine to uracil, and iii) propagation of the edit through DNA repair and/or cell division (Supplementary Figure 1A-B)^12^. DNA deamination occurs in a specific interval within the R-loop, called the editing window. As a result, cytosines other than the target residue are likely to be edited if they are present within the editing window^12^. While these so-called bystander edits are often problematic when the aim is to install or correct missense variants, CBE-based gene inactivation strategies, such as converting codons encoding amino acids into stop codons^13,14^ or mutating splice sites^15,16^, are generally not limited by the occurrence of bystander edits.

Predictable loss-of-function mutations introduced by base editing could circumvent some of the limitations associated with random mutations introduced by CRISPR nucleases, provided that they are installed with high efficiency and minimal undesired byproducts. However, several shortcomings and uncertainties exist. First, while CBEs and other base editors have been extensively optimised for expression, nuclear localization^17,18^ and activity of the deaminase domain^12,19^, loss-of-function experiments aiming to create biallelic gene knock-outs in all cells of a multicellular tissue or organism require particularly high editing rates, which remain difficult to attain with base editors in various systems^20–22^. This may at least partially be due to the fact that genome editing tools are typically developed and optimised in immortalised mammalian cell lines, which differ from many other systems relevant for research and biotechnology in parameters such as temperature and composition of DNA repair pathways. Second, while CBEs predominantly give rise to predictable C-to-T edits in the editing window, they are also known to cause random indel mutations through error-prone repair of abasic sites that arise from uracil removal by Uracil-N-Glycosylases (UNG)^10,12^, a process which can only partially be suppressed through UNG inhibitor domains (UGIs)^19,23,24^. Induction of such random indels limits the ability of CBEs to avoid undesirable editing products.

*Drosophila melanogaster* is one of the most widely used model organisms in biomedical research, largely due to the availability of sophisticated genetic tools^25^. Over the years, genetic research in *Drosophila* has uncovered numerous fundamental and evolutionary conserved principles governing the biology of multicellular animals^26–28^. Despite these efforts, functionally validated null alleles are still not available for the majority of genes encoded in the *Drosophila* genome^25^. In recent years, large collections of CRISPR gRNA lines have been generated that can be used in conjunction with Cas9 nuclease to induce targeted mutations^8,29–31^, but these suffer from the aforementioned limitations of random mutagenesis. So far, only a single study has reported on the use of a first-generation base editor in *Drosophila*, which was shown to induce C-to-T edits in less than half of the tested genomic target sites^21^. Whether optimized base editors can lead to more robust editing in this organism remains to be explored.

Here, we establish CBEs for gene inactivation in *Drosophila*, and uncover crucial parameters influencing the performance of these tools. While the efficiencies obtained with a derivative of the most commonly used APOBEC1 domain were constrained by the temperature range tolerated by *Drosophila*, a domain derived from an ectothermic species living in cold to temperate environments permitted robust base editing with remarkably high efficiency. Using this CBE in combination with gRNA multiplexing we observed near homogenous biallelic gene inactivation in almost all tested target genes *in vivo*. Moreover, we uncovered unusually high CBE product purity in *Drosophila*, which is likely attributable to the lack of a *UNG* gene in holometabola^32^. Our results highlight the usefulness of CBEs for gene inactivation in multicellular organisms, establish robust and highly efficient base editing tools for *Drosophila*, and reveal so far underappreciated parameters that influence the activity and purity of base editing systems.

## Results

### Generation of cytosine base editing tools

To explore CBE-mediated gene inactivation in *Drosophila*, we first constructed vectors for *in vivo* CBE expression. We utilised a Cas9^D10A^ coding sequence previously shown to be well tolerated in *Drosophila*^33^, linkers and bipartite nuclear localization signals (bpNLSs) from the optimised BE4max architecture^17^, as well as a UGI domain^10^ (Figure 1A). To explore the properties of two distinct deaminases, we constructed CBEs featuring recently laboratory evolved versions of the APOBEC1 protein from *Rattus norvegicus* and the CDA1 protein from *Petromyzon marinus*, respectively^19^ (henceforth referred to as *Drosophila* Cytosine Base Editor^evoAPOBEC1/evoCDA1^, dCBE^evoAPOBEC1^ and dCBE^evoCDA1^).

**Figure 1.**
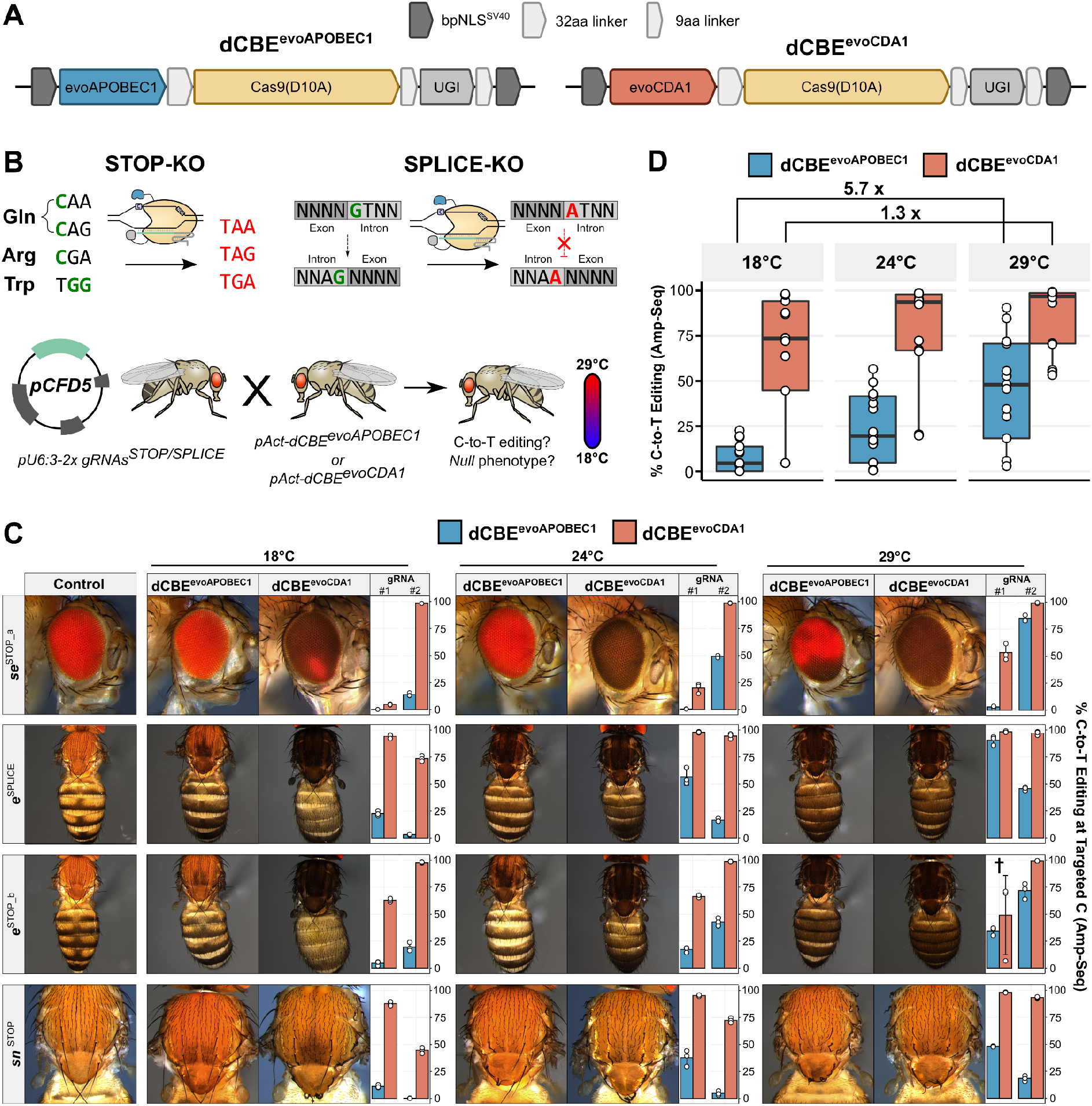
CBE-mediated gene inactivation across the thermal range tolerated by *Drosophila*. **(A)** Schematic representation of the base editor constructs used in this study. **(B)** Schematic representation of the experimental workflow. Pairs of gRNAs were designed to cause C-to-T edits either disrupting splice sites or generating *de novo* STOP codons. Males ubiquitously expressing the gRNAs were crossed to base editor-expressing virgins, and the offspring raised at different temperatures. **(C)** Representative images of female offspring are shown alongside the C-to-T editing rates at the targeted C residues, determined by Amp-Seq (n = 3, data presented as mean ± s.d. (bars with error bars) and individual measurements (points)). Whereas editing and phenotypic severity are strongly correlated with temperature for *pAct-dCBE^evoAPOBEC1^, pAct-dCBE^evoCDA1^* causes efficient gene inactivation across all conditions. The target site for the gRNA marked with a cross was found to harbour a floating polymorphism located two nucleotides upstream of the PAM, which was homozygous in one fly analysed for dCBE^evoCDA1^ at 29°C, giving rise to the outlying data point (see also Supplementary Figure 4B). **(D)** Effect of temperature on base editing efficiency. Each dot represents the mean (from n = 3) of a C residue located within positions 3-9 of any of the assessed target sites (8 gRNAs, 20 C residues). Horizontal line represents mean, hinges the first and third quartiles and whiskers 1.5 interquartile ranges. Fold-changes in average base editing efficiency between 18°C and 29°C are indicated.

For ubiquitous base editing, we cloned these editors downstream of the *act5c* promoter (*pAct-dCBE^evoAPOBEC1^* and *pAct-dCBE^evoCDA1^*, Supplementary Figure 2A) and generated transgenic animals harbouring a genomic insertions of these constructs on a defined landing site on the second chromosome. Such animals were viable, fertile and without any obvious phenotypic abnormalities and can be maintained as a stable stock under standard conditions (Supplementary Figure 2B). For conditional expression through the binary Gal4-UAS system (Supplementary Figure 2C), we generated animals transgenic for *pUAS-dCBE^evoAPOBEC1^* and *pUAS-dCBE^evoCDA1^*, respectively, which likewise were fertile and phenotypically normal. However, when crossing *pUAS-dCBE^evoAPOBEC1^* animals to *hh-Gal4* or *act-Gal4* driver lines to induce base editor expression, we did not obtain any viable offspring (Supplementary Figure 2D). In contrast, crosses with *pUAS-dCBE^evoCDA1^* produced viable offspring with both Gal4-drivers. However, *hh-Gal4* animals homozygous for *pUAS-dCBE^evoCDA1^* exhibited wing malformations indicative of reduced cell proliferation or excessive cell death, whereas animals with a single copy of the transgene did not.

Together, these findings reveal dose dependent toxicity with base editor constructs in *Drosophila*. While expression under direct control of the *act5C* promoter is well tolerated, expression under the Gal4-UAS system, which is known to lead to substantial overexpression, can lead to undesired side effects. Even though dose dependent toxicity *in vivo* of the Cas9 nuclease alone has been previously observed^8,34^, the effects of CBE overexpression observed here are more severe. Our finding that evoCDA1 is much better tolerated than evoAPOBEC1 suggests that the deaminase portion of the editors dictates tolerability and that CBE expression levels should be carefully controlled.

### Differential temperature sensitivity of evoAPOBEC1 and evoCDA1 base editors

Next, we set out to assess the efficiency of CBE-mediated gene inactivation. To facilitate both phenotypic and genotypic readouts, we initially targeted a set of non-essential genes with well-characterised, visible and recessive *null* mutant phenotypes (*ebony* (*e*), *singed* (*sn*), *forked* (*f*) and *sepia* (*se*)). To this end, we utilised the previously described *pCFD5* vector to ubiquitously express pairs of gRNAs^35^ designed to either mediate the conversion of codons coding for amino acids to STOP codons^13,14^ or to disrupt splice donor or acceptor sites^16^ at two independent positions in each gene (Figure 1B). To establish optimal conditions for base editing in *Drosophila*, we crossed males from four gRNA strains (one targeting *se*, two targeting *e* and one targeting *sn*) against *pAct-dCBE^evoAPOBEC1^* or *pAct-dCBE^evoCDA1^* virgins ubiquitously expressing either CBE, raised the animals at various temperatures within the range used to maintain *Drosophila* (18°C, 24°C and 29°C, Figure 1B) and assessed offspring phenotypically and by Illumina deep-sequencing of PCR amplicons (Amp-Seq) of the target sites.

We observed strong loss-of-function phenotypes when base editing was performed with dCBE^evoAPOBEC1^ at 29°C, with fully penetrant phenotypes observed with three gRNA strains and a mosaic phenotype with one strain (Figure 1C). However, phenotypes were attenuated when animals were raised at 24°C and nearly absent at 18°C (Figure 1C). In contrast, dCBE^evoCDA1^ mediated fully-penetrant phenotypes resembling heritable, homozygous loss-of-function alleles in all instances at 29°C and 24°C. Surprisingly, even at 18°C animals developed phenotypes indicative of biallelic gene disruption in most or all of their body.

We then used amplicon sequencing to sensitively quantify mutations at the target site of each gRNA. Consistent with our phenotypic observations, we found a strong correlation between temperature and C-to-T editing rates with dCBE^evoAPOBEC1^, but highly efficient editing at at least one target site across all conditions with dCBE^evoCDA1^ (Figure 1C). At the target residue C-to-T editing rates at 29°C with dCBE^evoAPOBEC1^ were 51.9±30% compared to 89.8±16.5% with dCBE^evoCDA1^, 29.7±20.3% compared to 82.4±26.5% at 24°C and 10.4±8.8% compared to 73.6±31.8% at 18°C. Summarising across all gRNAs and all C nucleotides within an editing window of positions 3-9 of the protospacer, base editing efficiency decreased an average of 5.7-fold from 29°C to 18°C with dCBE^evoAPOBEC1^, but only 1.3-fold with dCBE^evoCDA1^ (Figure 1D), demonstrating that activity of the evoCDA1 deaminase domain is much less temperature sensitive at that range.

Together, these findings demonstrate that the evoAPOBEC1 domain functions suboptimally at the lower temperatures preferred by *Drosophila* and other ectothermic organisms, whereas evoCDA1 permits efficient base editing within this broad temperature range.

### dCBE^evoCDA1^ permits highly efficient biallelic gene inactivation *in vivo*

To further explore whether dCBE^evoCDA1^ can mediate gene knock-out with high penetrance in somatic cells at a larger selection of target sites, we generated six additional pairs of gRNAs targeting the aforementioned non-essential genes as well as four pairs of gRNAs designed to create STOP codons in well-characterised essential genes (*wg*, *ct, evi* and *smo*).

In crosses targeting the non-essential genes at 29°C, *pAct-dCBE^evoCDA1^* led to fully penetrant loss-of-function phenotypes in all but one instance (Figure 2A and Supplementary Figure 3A). In the case of *f*^SPLICE_a^, where gRNAs target the splice acceptors of two small exons, we did not observe any phenotype indicating disruption of *f*. This could reflect inefficient base editing or result from biological compensation by exon skipping^15^ (Supplementary Figure 4A). Excluding the former possibility, we observed highly efficient (90.0±2.25% and 90.0±1.36%) C-to-T conversion at the target residue of both gRNAs present in *f*^SPLICE_a^ (Figure 2B). Together these results demonstrate that dCBE^evoCDA1^ mediates phenotypes with unusually high penetrance and robustness, but also highlights that the outcome of splice acceptor disruption is not always predictable (e.g. can for example lead to both intron retention or exon skipping), as has also previously been observed^15,16^.

**Figure 2.**
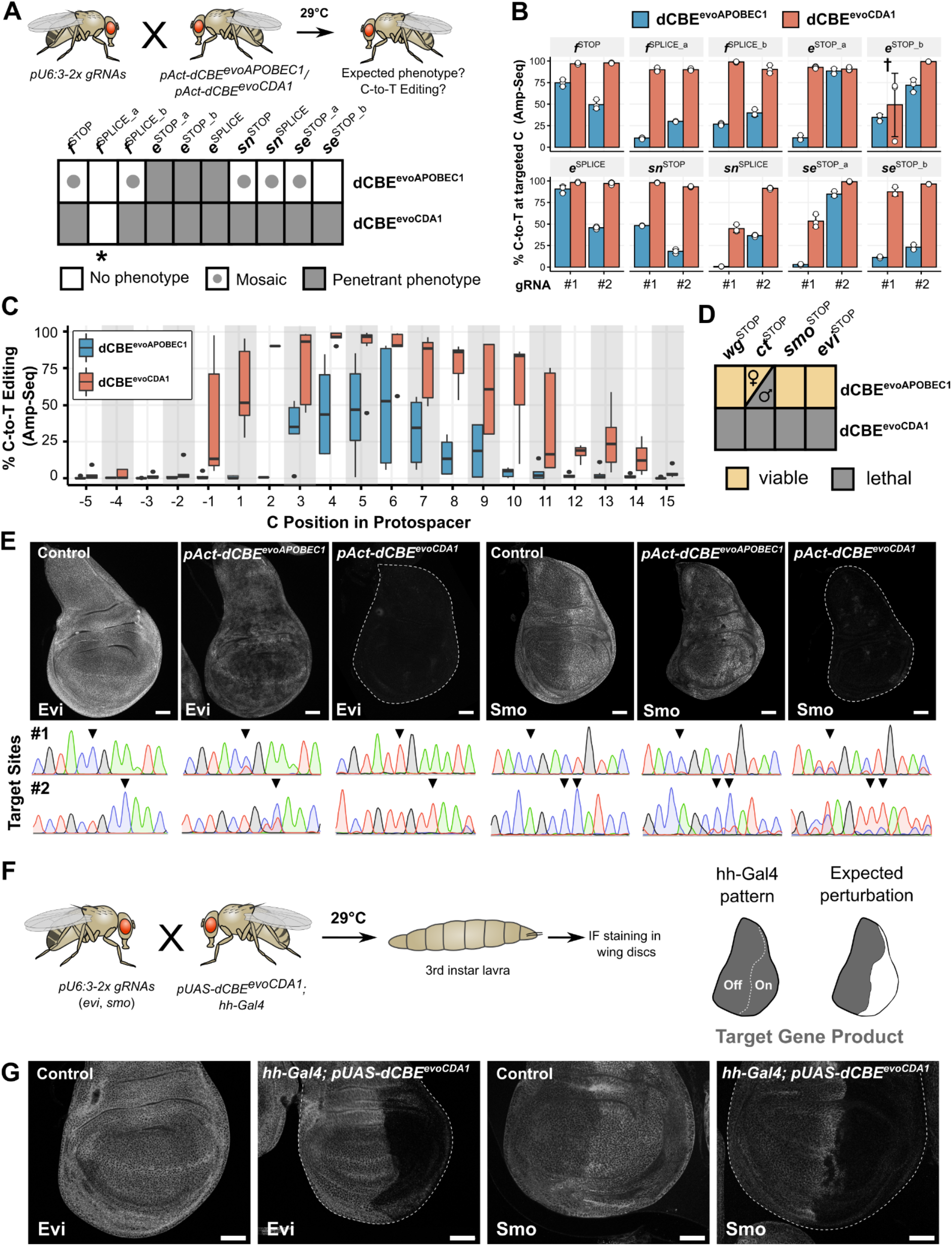
Highly efficient biallelic gene inactivation with dCBE^evoCDA1^. **(A)** Schematic representation of phenotypes obtained at 29°C with both base editors. While outcomes with *pAct-dCBE^evoAPOBEC1^* are heterogeneous, *pAct-dCBE^evoCDA1^* gives rise to penetrant phenotypes with all gRNA pairs except *f*^SPLICE_a^ (marked with an asterisk), which is likely the result of exon skipping caused by targeting of splice acceptor sites (see Supplementary Figure 3A for images of dCBE-induced phenotypes). **(B)** C-to-T editing rates obtained at the intended C residues, determined by Amp-Seq (n = 3, data presented as mean ± s.d. (bars with error bars) and individual measurements (points)). dCBE^evoCDA1^ achieves highly efficient C-to-T editing at one or both gRNAs for all tested pairs, including *f*^SPLICE_a^. The target site for the gRNA marked with a cross was found to harbour a floating polymorphism two nucleotides upstream of the PAM, which was homozygous in one fly analysed for dCBE^evoCDA1^ at 29°C, likely reducing base editing efficiency. **(C)** Editing efficiency of *pAct-dCBE^evoAPOBEC1^* and *pAct-dCBE^evoCDA1^* at 29°C plotted as a function of position in the protospacer (N = 20 target sites, individual data points are the mean from n = 3 Amp-Seq measurements for each target site). Horizontal line represents the mean, hinges the first and third quartiles and whiskers the 1.5 interquartile ranges. dCBE^evoCDA*1*^ has superior base editing efficiency and a broader editing window compared to dCBE^evoAPOBEC1^. **(D)** Mutagenesis of the essential genes *wg*, *ct, evi* and *smo*. Males ubiquitously expressing gRNA pairs were crossed to *pAct-dCBE^evoAPOBEC1^* or *pAct-dCBE^evoCDA1^* virgins. Animals were raised at 29°C and the viability of the offspring assessed. While dCBE^evoAPOBEC1^ crosses are mostly viable, dCBE^evoCDA1^ produces the expected outcome of lethality. **(E)** Immunofluorescence images and Sanger sequencing of 3rd instar larvae subjected to ubiquitous mutagenesis with the *evi*^STOP^ and *smo*^STOP^ gRNAs. While *pAct-dCBE^evoAPOBEC1^* gives rise to mosaics in the wing imaginal disc, *pAct-dCBE^evoCDA1^* leads to penetrant protein ablation in the entire tissue. Sanger chromatograms exhibit incomplete C-to-T editing with *pAct-dCBE^evoAPOBEC1^* but efficient conversion with *pAct-dCBE^evoCDA1^.* Scale bars represent 50 μm. Triangles in the sequencing chromatograms highlight the target C residue. **(F)** Schematic representation of tissue-specific mutagenesis experiments in the wing imaginal disc using the hh-Gal4 driver to drive dCBE^evoCDA1^ expression in the posterior disc compartment. **(G)** Representative immunofluorescence images of wing imaginal discs from offspring of *hh-Gal4; pUAS-dCBE^evoCDA1^* animals crossed against *evi*^STOP^ or *smo*^STOP^ gRNAs and raised at 29°C are shown. Evi and Smo protein loss is detected in the posterior compartment, with no signs of ectopic mutagenesis in the anterior compartment. White arrows indicate instances of morphological abnormalities. Scale bars represent 50 μm.

In contrast to our results with dCBE^evoCDA1^, phenotypic outcomes resulting from editing with dCBE^evoAPOBEC1^ were much more heterogeneous. We observed fully penetrant phenotypes in three instances, mosaic phenotypes with five gRNA pairs and no detectable phenotype in two conditions (Figure 2A and Supplementary Figure 3B). Sequencing of the gRNA target sites confirmed highly efficient C-to-T conversion of the target deoxycytidine with dCBE^evoCDA1^, but large variation in base editing efficiency at the target base as the underlying cause of variation in phenotypic penetrance with dCBE^evoAPOBEC1^ (Figure 2B). We tested if a publicly available machine-learning model could predict which gRNAs support efficient editing with dCBE^evoAPOBEC1^, but found only a modest and not statistically significant correlation between the Z-score predicted by the model^36^ and the observed fraction of base-edited alleles at each target site (Pearson *R* = 0.34, Supplementary Figure 5).

To determine the average base editing efficiency across the base editing window at 29°C with both editors, we calculated the averaged C-to-T conversion rates as a function of position in the protospacer. In line with previous reports for these deaminases^19^, dCBE^evoAPOBEC1^ and dCBE^evoCDA1^ exhibited editing windows of different size, spanning positions 3-9 and −1-14, respectively (Figure 2C, defined as > 10% median C-to-T editing efficiency). While dCBE^evoAPOBEC1^ achieved a maximal mean editing efficiency of 45.9%-53.5% at positions 4-6 of the protospacer, dCBE^evoCDA1^ enabled highly robust C-to-T conversion of 88.3%-96.3% at the same positions of the target sites.

Next, we crossed the *pAct-dCBE* constructs to gRNA pairs targeting a set of essential genes (*wg*, *smo*, *evi* and *ct*). While *dCBE^evoCDA1^/gRNA* animals died during development, the known loss-of-function phenotype of these target genes, crosses with dCBE^evoAPOBEC1^ gave rise to viable offspring with wing malformations (Figure 2D and Supplementary Figure 6), indicating inefficient mutagenesis. To confirm this, we stained wing imaginal discs from 3rd instar larvae from crosses targeting *evi* and *smo* for endogenous Evi and Smo protein and assayed the status of each target site by Sanger sequencing. While dCBE^evoAPOBEC1^ caused incomplete loss of protein expression in the wing disc and led to limited editing of the target residues, dCBE^evoCDA1^ mediated almost complete ablation of Evi and Smo in all cells of the tissue and caused efficient C-to-T editing at the targeted C residues (Figure 2E).

A useful strategy to circumvent lethality when targeting essential genes is to restrict mutagenesis to the target tissue of interest. To investigate if base editing permits the tissue-selective inactivation of essential gene targets, we performed conditional mutagenesis in the wing imaginal disc by utilising the *hh-Gal4* driver, which is selectively expressed in the posterior compartment of this tissue (Figure 2F). We crossed *pUAS-dCBE^evoCDA1^; hh-Gal4* virgins to males expressing gRNAs targeted against either *evi* or *smo*, raised the animals at 29°C and performed immunofluorescence analysis on wing imaginal discs from 3rd instar larvae (Figure 2F). Compared to control discs, dCBE^evoCDA1^ expression caused some morphological abnormalities (Supplementary 7, white arrows), in line with our previous observations of overexpression-related toxicity. gRNA pairs targeting *evi* or *smo* resulted in ablation of the respective gene products in the posterior compartment only, with no signs of residual gene product in the targeted area nor evidence of ectopic mutagenesis in the non-targeted compartment (Figure 2G).

In summary, these results establish that dCBE^evoCDA1^ in conjunction with two gRNAs enables highly efficient biallelic gene inactivation *in vivo*, with practically no unedited or function-retaining alleles, in both ubiquitous as well as tissue-specific settings.

### Creating heritable alleles with base editing in the *Drosophila* germline

While a previous study established base editing with a first-generation CBE in *Drosophila* somatic cells^21^, germline transmission of base edits has thus far not been described. We reasoned that dCBE^evoAPOBEC1^ might be the preferred editor for germline transmission applications, as one typically screens several animals of the offspring, meaning intermediate editing efficiencies are sufficient, and a narrower editing window makes it more likely to recover precise edits at the target C without bystander edits.

We first performed a genetic complementation assay by crossing *pAct-dCBE^evoAPOBEC1^;pCFD5-2xgRNA* males raised at 29°C to virgins harbouring homozygous null alleles in the targeted genes, and assessed the percentage of offspring inheriting two loss-of-function alleles and thus exhibiting the respective null mutant phenotype (Figure 3A). For all four assayed gRNA lines, we detected germline transmission rates that were in line with the somatic editing efficiencies we had observed previously (84%, 95%, 70% and 75% respectively, Figure 3B). To confirm that phenotypes were caused by the intended gene edits and to exclude unintended indel byproducts, which have been previously shown to occur with CBEs^10,19^, we performed Sanger sequencing of the *e*^STOP_b^ and *e*^SPLICE^ target sites in phenotypic offspring to assess the identity of the transmitted edits. In every instance (*N = 22*) we detected at least one base edit at one of the two target sites and did not observe any indels (Figure 3C-D), confirming that our assay accurately reflected the level of base editing in the germline. At the first target site of *e*^STOP_b^ we recovered different base edited alleles, some exclusively with C-to-T editing at the target C and others with an additional edit at another residue in the target window (Figure 3D), demonstrating that the intermediate efficiency of dCBE^evoAPOBEC1^ can be used to recover different alleles in the same cross. Together these results demonstrate efficient base editing in the *Drosophila* germline to create heritable alleles.

**Figure 3.**
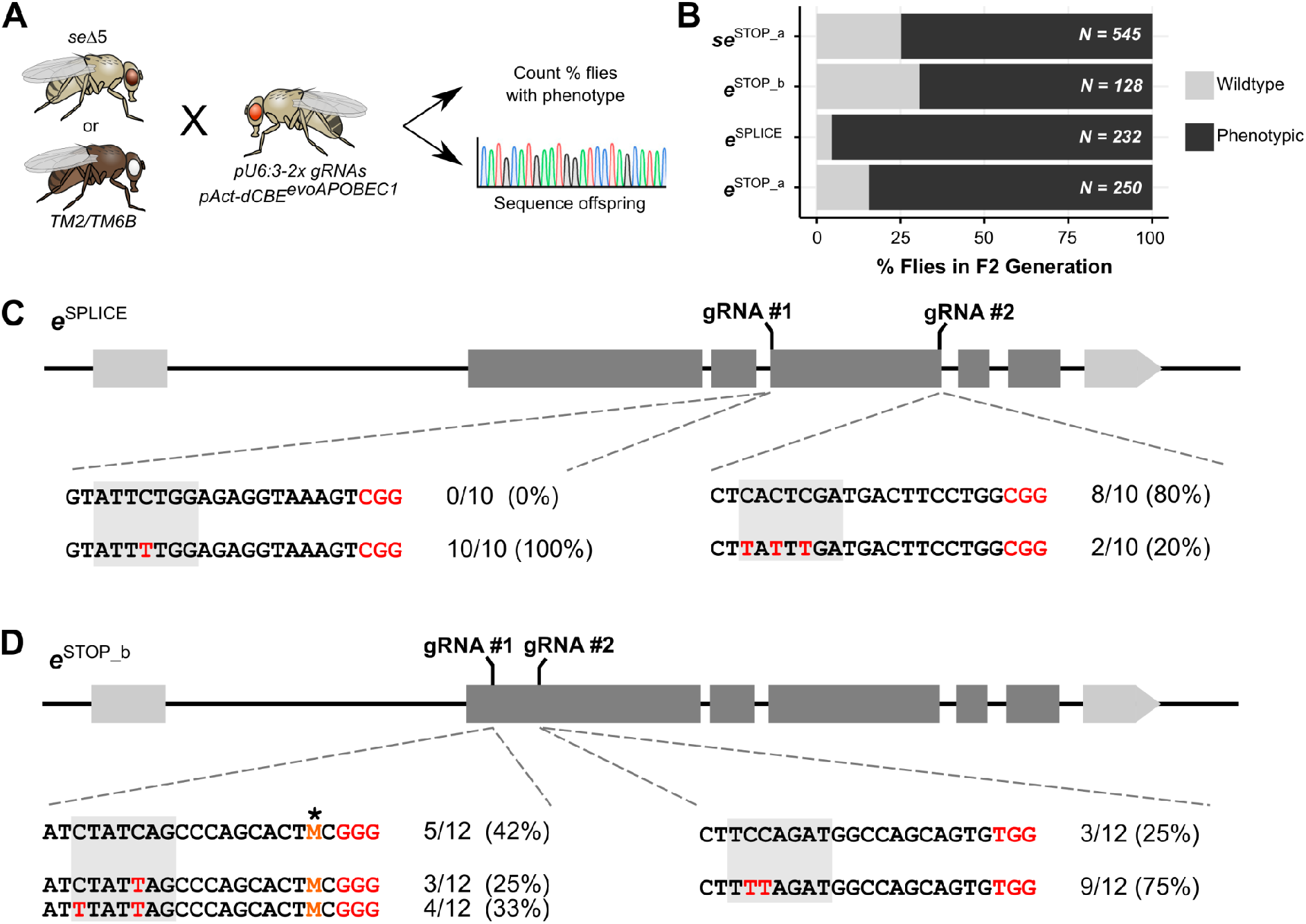
Germline transmission of base-edited alleles. **(A)** Schematic of the genetic complementation assay for germline transmission. F1 males harbouring both genome editing transgenes were crossed to homozygous mutant flies (*seΔ5* or *TM2/TM6B*, for transmission of *se* or *e* alleles, respectively) and transmission of base-edited alleles from the male parent determined by phenotypic assessment and Sanger sequencing. **(B)** Germline transmission of base-edited alleles as percentage of F2 flies exhibiting the respective phenotype (assessed from a total of n = 3 independent crosses). Transmission rates for each gRNA strain are reflective of the editing rates observed with Amp-Seq. **(C-D)** Alleles observed in F2 offspring with the *e*^SPLICE^ and *e*^STOP_b^ gRNAs, respectively. Shown are the protospacer (black) and PAM (red) of the target sites, as well as the editing window in grey. Types and frequency of the observed alleles are shown. Only reference or alleles carrying C-to-T edits are observed. The position carrying a SNP is shown in orange and highlighted with an asterisk (where M is A or C).

### Base editing by-product formation is exceptionally rare in *Drosophila*

Intrigued by the complete absence of indel byproducts in our germline transmission experiment, we investigated our more sensitive Amp-Seq dataset for the presence of indel byproducts in flies mutagenized with *pAct-dCBE^evoAPOBEC1^* or *pAct-dCBE^evoCDA1^* at 29°C.

The average indel frequencies with dCBE^evoAPOBEC1^ and dCBE^evoCDA1^ were 0.5% and 2.3%, respectively, which is substantially below the indel rates reported using the same deaminases in human cells (8.9% and 12.4%, respectively, Figure 4A)^19^. Likewise, C-to-G or C-to-A edits, another common type of undesired CBE by-products, were virtually absent at the vast majority of target sites (Figure 4B), despite the very high C-to-T editing rates (see Figure 2B). The indels we did observe were highly diverse and frequently comprised the editing window (Supplementary Figure 8A), suggesting deaminase rather than Cas9 nickase activity as causative for these edits. Supporting this hypothesis, target sites with four C residues in their editing window tended to show higher indel formation rates than sites with fewer C nucleotides (Supplementary Figure 8B).

**Figure 4.**
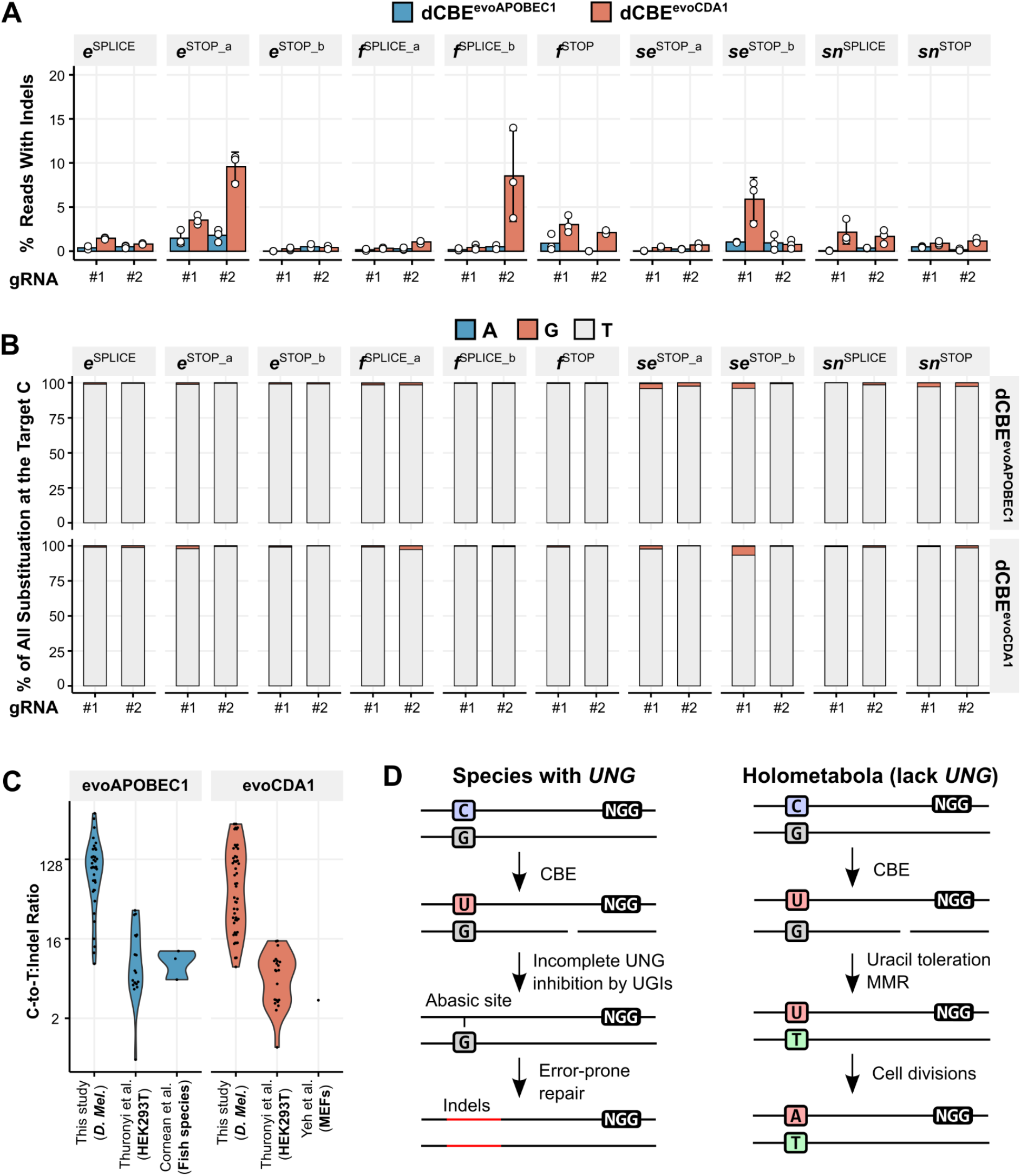
Analysis of by-products caused by cytosine base editing. **(A)** Percentage of Amp-Seq reads containing indels at 29°C (n = 3, data presented as mean ± s.d. and individual measurements). With some exceptions, indel formation is generally low (<2% at most target sites) and slightly higher with dCBE^evoCDA1^ compared to dCBE^evoAPOBEC1^. **(B)** Frequency of different types of base substitution observed at the target C residues. With both deaminases and across the vast majority of target sites, virtually all observed substitutions are C-to-T, with only a few target sites exhibiting low levels of C-to-G conversion (data presented as the mean frequency of reads exhibiting the respective substitution out of all reads harbouring a non-C nucleotide at the target residue). **(C)** C-to-T over indel ratio observed with evoAPOBEC1 and evoCDA1 in this and other studies. Shown are the ratios for all C-residues at positions 3-10 of the target sites that exhibit at least 5% C-to-T conversion within the respective datasets. Compared to observations in other systems, the C-to-T:indel ratios are drastically elevated in *Drosophila*. **(D)** Proposed mechanism for reduced indel formation in *Drosophila*. In species harbouring a *UNG* gene, incomplete inhibition of the enzyme by UGI domains results in formation of abasic sites, which are repaired in an error-prone fashion. In *Drosophila* and presumably other holometabola, Uracil is tolerated in genomic DNA, favouring mismatch repair and resolution to a desired C-to-T edit.

To control for potential differences in the overall base editing activity between our experiments and other studies, we calculated the C-to-T:indel ratio for C residues with at least 5% base conversion at any given target site. With both dCBE^evoAPOBEC1^ and dCBE^evoCDA1^ most C residues exhibited C-to-T:indel ratios between 30 and 300 (Figure 4C). In contrast, other studies using evoAPOBEC1 and evoCDA1 in immortalised human cells^19^, mouse embryonic fibroblasts^37^, medaka or zebrafish^38^ showed C-to-T:indel ratios between 5 and 25 for evoAPOBEC1 and between 2 and 16 for evoCDA1 (Figure 4C). This reveals that base editing-induced indel formation is exceptionally rare in *Drosophila*. Intriguingly, *Drosophila* and other Holometabola lack a *UNG* gene^32^, suggesting a mechanism where Uracil toleration leads to reduced indel formation and increased installation of desired C-to-T edits in these species (Figure 4D), which is in line with a previous study showing substantially improved CBE product purity in human *UNG* knockout cells^23^. The few observed indels might be caused by thymine DNA-glycosylase (TDG), which is present in *Drosophila* and was previously shown to be active on U:G mismatches *in vitro*^39^.

In summary, these analyses show that aside from achieving high C-to-T conversion rates, CBEs in *Drosophila* also exhibit remarkably elevated product purity and suggest that current strategies for *UNG* inhibition in other species are only partially successful.

## Discussion

In this study, we establish efficient CBEs in *Drosophila* for gene inactivation through C-to-T editing. The system is composed of genetically encoded CBEs and gRNAs to create premature stop codons or to disrupt splice sites. By combining a base editor featuring the evoCDA1 deaminase domain with high activity at temperatures below 30°C and multiplexing of two gRNAs targeting the same gene, biallelic knock-out alleles can be installed in most or all cells of the animal. Furthermore, the high product purity attainable with base editing in *Drosophila* minimises the generation of function-retaining alleles, further contributing to the very high efficiency of this system.

Efficiency of gene disruption experiments with CRISPR nucleases in bulk cell populations and entire organisms are usually curbed by limited gRNA activity and the generation of function-retaining mutations at the target site^4,7^. Multiplex base editing presents a broadly applicable alternative to precisely install loss-of-function alleles. The generation of stop codons is hereby more widely applicable, as typically there are many more positions in a gene where it is possible to convert codons to stop codons than there are targetable splice sites, in particular in organisms with compact genomes^13,14^. For example, we were not able to design SPLICE gRNAs against the small *se* gene. Moreover, when targeting splice sites, splice acceptors of small exons should be avoided given the tendency to result in exon skipping, as previously described^15,16^ and also evidenced by the *f*^SPLICE_a^ gRNAs in our study. In contrast, premature stop codons are expected to have severe functional consequences on gene function, in particular when placed in the 5’ portion of the gene, where they are also more likely to cause nonsense mediated mRNA decay^40,41^. However, the possibility of stop codon read-through and functionality of polypeptides encoded 5’ of the first base edited allele should be considered when designing and analysing such experiments. For both strategies, computational tools are available for the identification of suitable gRNAs^13,16^ (see also Methods). Notably, such gRNAs can in many cases also be used in combination with existing Cas9 nucleases to create mutations by error-prone double strand break repair, should both NHEJ and base-editing mediated gene disruption be of interest.

Our work reveals the crucial role of temperature in determining base editing efficiency with different CBEs. While the evoAPOBEC1 deaminase domain shows rapidly declining activity at temperatures below 30°C, evoCDA1 is better adapted to this temperature range, likely reflecting the fact that these deaminase domains originate from an endothermic and ectothermic organism, respectively. While the fact that different enzymes are optimised for different temperatures is generally recognized, temperature is rarely a tested parameter during the development of novel gene editing tools. As a result, the optimal temperature range for most genome editors is not known. Prior work has revealed that variants of the Cas12a nuclease derived from different bacterial species vary dramatically in their activity at lower temperature^42,43^. Our study extends this observation to deaminases used for precision genome engineering. Outside of applications in mammals, many of the most important uses of CRISPR genome engineering take place in organisms that do not tolerate longer term exposure to temperatures of 37°C. For example, important disease vectors, such as mosquitos of the genus *Aedes* that transmit malaria, dengue and other diseases, are optimally adapted to temperatures between 20°C and 30°C, with an upper limit of 34°C^44^. Likewise, the optimal growth temperatures for the majority of crop plants is below 30°C^45^. It is therefore important to develop genome engineering tools optimised to different climates to realise the full potential of CRISPR technology.

One surprising finding of our study is the fact that base editing in *Drosophila* leads to much fewer undesired byproducts in the form of indels or C-to-G or C-to-A edits at the gRNA target site compared to many other species. This is likely due to system specific differences in DNA repair. Indel byproducts during cytosine base editing mostly arise from error-prone repair of abasic sites that are generated by the UNG enzyme^10,23^. To limit such events, CBEs are typically linked to one or more UNG-inhibition (UGI) domains^23,24^. *Drosophila* and other holometabolous insects are naturally missing a *UNG* gene^32^ and thus mimic human *UNG* knockout cells, which had previously been shown to essentially eliminate indels and C-to-G/A editing^23^. The fact that we did observe some indel formation, in particular with the highly active evoCDA1 domain, suggests that additional DNA repair enzymes recognize intermediate base editing products and contribute to the formation of undesired editing outcomes. For example thymine DNA-glycosylase (TDG) has previously been shown to act on U:G mismatches^39^, suggesting additional targets to increase base editing precision.

The data presented here also highlight organismal toxicity as a potential limitation of current base editing technology. While base editing does not induce genotoxic DNA double strand breaks, it does rely on the expression of constitutively active deaminase domains, which on their own act indiscriminately on single stranded DNA and in some cases RNA^46,47^. Specificity of such systems relies on the ability to concentrate an active base editor at the target site through RNA-guided DNA binding of Cas9, while limiting base editor concentration in the rest of the cell. Indeed, a previous study has shown that limiting CBE expression to the level of housekeeping genes abrogates the Cas9-independent DNA and RNA off-targeting observed with transfection mediated overexpression^48^. In line with this, we find that CBEs expressed directly from a housekeeping promoter (*act5c*) are well tolerated in *Drosophila*, while overexpression with the binary expression system Gal4-UAS gives rise to severe side effects. As the expression levels achieved with the *act5c* promoter already mediate highly efficient base editing, it should be possible to reduce expression by the Gal4-UAS system to minimise toxicity with limited or no trade-off in efficiency, as previously demonstrated for Cas9 and a DNA methyltransferase^8,49^. Interestingly, toxicity arising from CBE overexpression is also dependent on the deaminase domain. While dCBE^evoAPOBEC1^ resulted in lethality when expressed under the control of an *act*- or *hh-Gal4* driver, dCBE^evoCDA1^ was better tolerated under the same conditions, despite the higher activity of this editor. It is tempting to speculate that this at least in part is due to the fact that APOBEC1, but not CDA1, exhibits RNA off-targeting in human cells^46,50^. In the future, the use of engineered deaminase domains with reduced Cas9-independent off-targeting^50,51^ might further reduce the toxicity of base editors in *Drosophila* and other species. Likewise, recently described TadA-based CBEs have been shown to minimise the occurrence of DNA and RNA off-targeting^52,53^.

In summary, our study describes highly efficient base editing tools for the installation of loss-of-function alleles in somatic cells or the germline of *Drosophila*. These systems will be useful for the efficient establishment of mutant fly lines or testing of gene function in the F1 generation. Base editor-mediated gene inactivation will be particularly advantageous in scenarios where function-retaining indels are problematic, knowledge of the type of edit is desired, deleterious consequences of DSBs are particularly problematic or when PCR bias would obscure sequencing-based analysis of target sites of different length. Our work also highlights several avenues to further improve CRISPR technology, such as optimization of temperature range and expression levels and leveraging the natural diversity in DNA repair between different systems. Thus, the tools and strategies described herein should have broad usefulness for loss-of-function experiments in *Drosophila* and other multicellular organisms and inform future developments of CRISPR technology.

## Methods

### Plasmid construction

A list of oligonucleotides and gBlock dsDNA fragments used in this study is provided in Supplementary Table 1. PCRs were performed using Q5 Hot-start 2× master mix (New England Biolabs). Restriction digests were conducted for 2 hours at 37°C. The linearized plasmid backbones and PCR products were analysed on agarose gels and purified using the QIAquick gel extraction kit (Qiagen). Assembly of DNA fragments was either performed with isothermal assembly or Goldengate-like reactions (see details on individual plasmids below). Reactions were transformed into Stellar competent cells (Takara) and plated on lysogeny broth (LB) agar plates supplemented with 100 μg/mL carbenicillin. Individual colonies were grown in LB broth supplemented with 100 μg/mL carbenicillin for 16 hours at 37°C and 180 rpm. The plasmid DNA was extracted using the QIAprep spin miniprep kit (Qiagen) and newly inserted sequences verified by Sanger sequencing.

#### pAct-dCBE^evoAPOBEC1^

DNA encoding the N- and C-terminal CBE components were ordered as gBlocks (Integrated DNA Technologies) and PCR amplified. The Cas9(D10A) coding sequence was generated through PCR amplification from plasmid *pAct-Cas9*^33^ using primers *Cas9D10Afwd/Cas9D10Arev*. The purified PCR products were assembled into EcoRI-HF/XhoI-digested *pAct-Cas9*^33^ by In-Fusion cloning (Takara).

#### pAct-dCBE^evoCDA1^

A gBlock encoding the evoCDA1 domain and Cas9 N-terminus until the NdeI restriction site in *pAct-dCBE^evoAPOBEC1^* was amplified using primers *oRMD204/oRMD205*. A second PCR was conducted on *pAct-dCBE^evoAPOBEC1^* with primers *oRMD158/oRMD206* to generate a fragment with overlaps to the upstream NdeI site and 5’ end of the evoCDA1 domain. The PCR fragments were assembled into NdeI-digested *pAct-dCBE^evoAPOBEC1^* using NEBuilder^®^ HiFi DNA assembly (New England Biolabs).

#### pUAS-dCBE^evoAPOBEC1^

The dCBE^evoAPOBEC1^ coding sequence was amplified from the *pAct-dCBE^evoAPOBEC1^* plasmid using primers *oRMD134/oRMD135* and the purified product assembled into EcoRI-HF/XbaI-digested *pUASTattB* backbone^54^ using NEBuilder^®^ HiFi DNA assembly (New England Biolabs).

#### pUAS-dCBE^evoCDA1^

The dCBE^evoCDA1^ coding sequence was amplified from the *pAct-dCBE^evoCDA1^* plasmid using primers *oRMD134/oRMD135* and the purified product assembled into EcoRI-HF/XbaI-digested *pUASTattB* backbone^54^ using NEBuilder^®^ HiFi DNA assembly (New England Biolabs).

#### pCFD5-2× gRNAs

2× gRNA vectors were constructed as previously described^8^. Forward and reverse primers encoding BbsI handles and the spacer sequences of gRNAs #1 and #2 were used in PCRs on plasmid *pCFD6*^35^. The purified reaction products were pooled at equal molarity and assembled into *pCFD5* in a Goldengate reaction using BbsI-HF and T4 Ligase (both New England Biolabs).

### CBE gRNA Design

The iSTOP R package^13^ and SpliceR webtool^16^ (https://moriaritylab.shinyapps.io/splicer/) were used to identify potential STOP and SPLICE gRNAs, respectively. Target sites for experimental interrogation were then selected based on protospacer composition (40-60% GC, balanced nucleotide representation), relative location in the coding sequence of the gene (first half), and position of the target residue within the editing window (5-7). In cases when no such two target sites were available, gRNAs at more 3’ regions in the gene and target residues at more sidewards position of the editing window (4-9) were chosen. Spacer sequences of all gRNAs used in this study are provided in Supplementary Table 2.

### *Drosophila* Strains and Handling

#### General Husbandry

All *Drosophila* strains generated and/or used in this study are listed in Supplementary Table 3. Flies were kept at 50 ± 10% humidity in a 12h light/12h dark cycle. The temperatures at which individual experiments were conducted are indicated in the figures. Temperature was regularly checked with thermometers.

#### Genomic DNA preparation

Material from individual flies (generally heads, if not otherwise indicated) were collected in 20 μL squishing buffer (10 mM Tris·HCl pH8, 1 mM EDTA, 25 mM NaCl, 200 μg/mL Proteinase K). For genomic DNA extraction from larvae, a dissection was performed to remove gut and fat tissue. The tissue was then disrupted in a Bead Ruptor (Biovendis) for 20 seconds at 25 Hz. The samples were incubated for 30 minutes at 37°C for proteinase K digestion and heat inactivated at 95°C for 3 minutes.

#### Transgenesis and genotyping

Transgenesis was conducted using the PhiC31/attP/attB system^54^. Prior to injections, plasmid DNA was purified using the QIAquick PCR purification kit (Qiagen) and normalised to 250 ng/ul in nuclease-free water. For pooled injections, an equimolar ratio of all plasmids was pooled. Microinjections were performed into *y[1] M{vas-int.Dm}ZH-2A w[*]; (P{y[*+*t7.7]CaryP}attP40)* blastoderm embryos for 2nd chromosomal integration and on *y[1] M{vas-int.Dm}ZH-2A w[*]; (P{y*+-*attP-3B}VK00033)* embryos for 3rd chromosomal integrations following standard procedures. The embryos were then raised at 18°C until reaching the larval stage (approximately 48 hours). The animals were then transferred to 24°C and raised to adulthood and individual flies crossed to *P{ry[*+*t7.2]* = *hsFLP}1, y[1] w[1118]; Sp/CyO-GFP* or *w[*]; ;Sb/TM6* balancer flies for 2nd and 3rd chromosomal integrations, respectively. Transgenic offspring were identified by eye colour and then crossed against appropriate balancer flies to generate a stable genetic stock.

If multiple gRNA plasmids were injected in a pool, transgenic adults were subjected to genotyping after having produced sufficient numbers of offspring. To this end, 2 μL of genomic DNA (gDNA) was utilised in PCRs with primers *pCFD8genofwd5*/*pCFD8genorev6*, and the purified PCR product subjected to Sanger sequencing with primer *pCFD8seqonly*.

### Germline Transmission of Base Edits

*P{ry[*+*t7.2]* = *hsFLP}1, y[1] w[1118]; pAct-dCBE^evoAPOBEC1^/+;pU6:3-gRNAs/+* males obtained in the F1 generation were crossed to *w[*]; Bl/CyO; TM2/TM6B* virgins for gRNAs targeting *ebony* and *seΔ5* virgins for gRNAs targeting *sepia*. The percentage of phenotypically *ebony* or *sepia* flies was then determined in the F2 generation. To exclude F2 animals that inherited both genome editing transgenes from their male parent, we only assessed animals lacking at least one of the two transgenes (determined by fainter eye colour). For transmission of *sepia* alleles, this exclusion was achieved based on the presence of somatic *sepia* mosaics that were observed when an F2 animal inherited both genome editing components and a functional *sepia* allele in their germline. For molecular determination of the transmitted edits, gDNA was extracted from F2 animals and the target loci amplified by PCR. The purified reaction products were submitted to Sanger sequencing and the transmitted edits identified by visual inspection of the sequencing chromatograms.

### Sanger sequencing of base editing target sites

Genomic loci were amplified using primers located at least 50 nucleotides 5’ and 3’ of the target sites of a given gRNA strain. The reactions were purified with paramagnetic beads and submitted for Sanger sequencing

### Quantification of Genome Editing Outcomes by Deep Amplicon Sequencing (Amp-Seq)

#### Library Preparation

Targeted deep sequencing of CRISPR edits was performed using a dual PCR-based strategy similar to previously described^55^.

Briefly, in a first PCR, genomic target sites were PCR-amplified (Q5® Hot Start High-Fidelity DNA Polymerase, New England Biolabs) using primers encoding Illumina adapter sequences (Supplementary Table 4). Reaction clean-up and size selection was performed using paramagnetic beads prepared as previously described^56^. The concentrations of the purified reaction products were determined on a DropSense 96 (Trinean) and approximately equal amounts of different genomic amplicons then pooled and subsequently diluted in nuclease-free water by a factor of 10. A second PCR was then performed on the diluted amplicon pools with primers adding Illumina indexes and P5/P7 sequences (Supplementary Table 4). The reactions were purified and size-selected using paramagnetic beads, and then quantified on a DropSense 96 (Trinean). Equal amounts of all libraries were combined into a final multiplex, which was quantified with a Qubit 1.0 fluorometer (Thermo Fisher Scientific) and then sequenced on an Illumina MiSeq (150-bp paired-end reads) by the High-Throughput Sequencing Unit of the Genomics and Proteomics Core Facility (German Cancer Research Center, DKFZ).

#### Data Analysis

For data analysis, CRISPResso2^57^ was run in pooled mode with the following parameters: -w 20 --cleavage_offset −10 --base_editor_output --min_reads_to_use_region 100. The sequences of the genomic amplicons are provided in Supplementary Table 5. Nucleotide conversion rates at individual nucleotides were extracted from the Quantification_window_nucleotide_percentage_table.txt files. The number of reads containing insertions or deletions was extracted from the CRISPResso_quantification_of_editing_frequency.txt files, and the percentage of reads containing indels then calculated as [# reads containing indels]/[# reads aligned]. For amplicons containing SNPs in their quantification window, CRISPResso2 was run with both alleles to avoid false classification of SNPs as NHEJ events. Indel sizes were extracted from the Indel_histogram.txt files.

### CBE Efficiency Prediction with BE-HIVE

BE-Hive^36^ (https://www.crisprbehive.design/) was run in single mode and provided with 20 nucleotide sequence contexts upstream and downstream of the spacers (i.e. a total of 60 nucleotides of the genomic target region). ‘evoAPOBEC1, mES’ was selected as base editor and cell type setting. The Z-scores predicted by the model were correlated with the observed base editing rates.

### Immunohistochemistry

Third instar wandering stage larvae were dissected in ice cold PBS using standard procedures. Larvae were fixed in 4% paraformaldehyde for 25 min at room temperature. Larvae were washed three times 15 min with PBT (1× PBS with 0.3% Triton-X100) and blocked for 30 min in PBT containing 1% normal goat serum. Samples were incubated with first antibodies (rabbit anti-Evi^58^ 1:500), mouse anti-wg (DSHB 4D4, 1:200), mouse anti-smo (DSHB, 1:50), mouse anti-ct (DSHB, 1:20)) overnight at 4°C. Following three 15 min washes with PBT samples, were incubated with secondary antibodies (Invitrogen, goat anti-rabbit or goat anti-mouse conjugated to Alexa fluorophores, 1:400 in PBT containing DAPI) for 2 hours at room temperature. Larvae were washed three times for 15 min at room temperature and wing imaginal discs were mounted in Vetrashield (Biozol).

### Image Acquisition and Processing

Microscopy images were acquired on a Leica LSM SP8 confocal microscope with a 40× NA1,3 Oil immersion lens in sequential scanning mode. Samples of the same experiment were imaged in the same session. Images were processed using Fiji.

## Acknowledgments

We would like to thank Ann-Christin Michalsen, Alexander Kremer, Martha Buhmann, Eva Roßmanith and Amélie Pöltl for help and discussions. We thank Benjamin Kleinstiver (MGH and Harvard University) for advice on Amp-Seq. We thank the High-Throughput Sequencing and Light Microscopy core facilities of DKFZ for support. This work was in part funded by grants from Deutsche Forschungsgemeinschaft (SFB1324) and the European Research Council (ERC Synergy Project DECODE).

## Author contributions

RMD, MB and FP conceived the project. RMD and FP designed and RMD performed experiments and RMD and FP analyzed the data. RMD and FP wrote and MB edited the manuscript. MB acquired funding and MB and FP co-supervised the work.

## Material availability

Material is available upon request from the corresponding authors. Plasmid encoding base editors will be available from Addgene (https://addgene.org, ID’s 195514, 195515) and the European Plasmid Repository (https://plasmids.eu). High-throughput sequencing data has been deposited at the EMBL-EBI European Nucleotide Archive project accession PRJEB58121.

## Supplementary Tables

**Supplementary Table 1.**
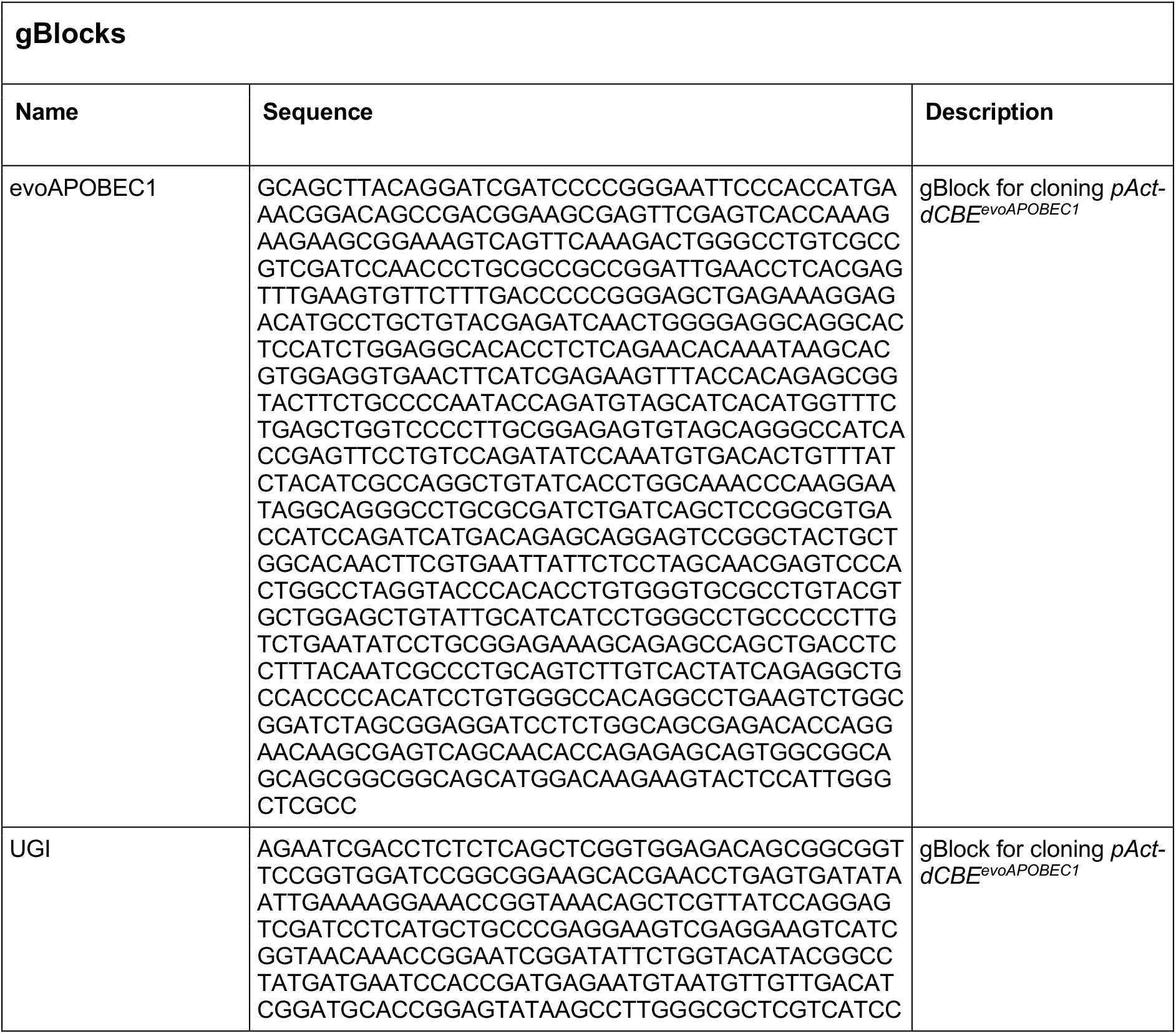
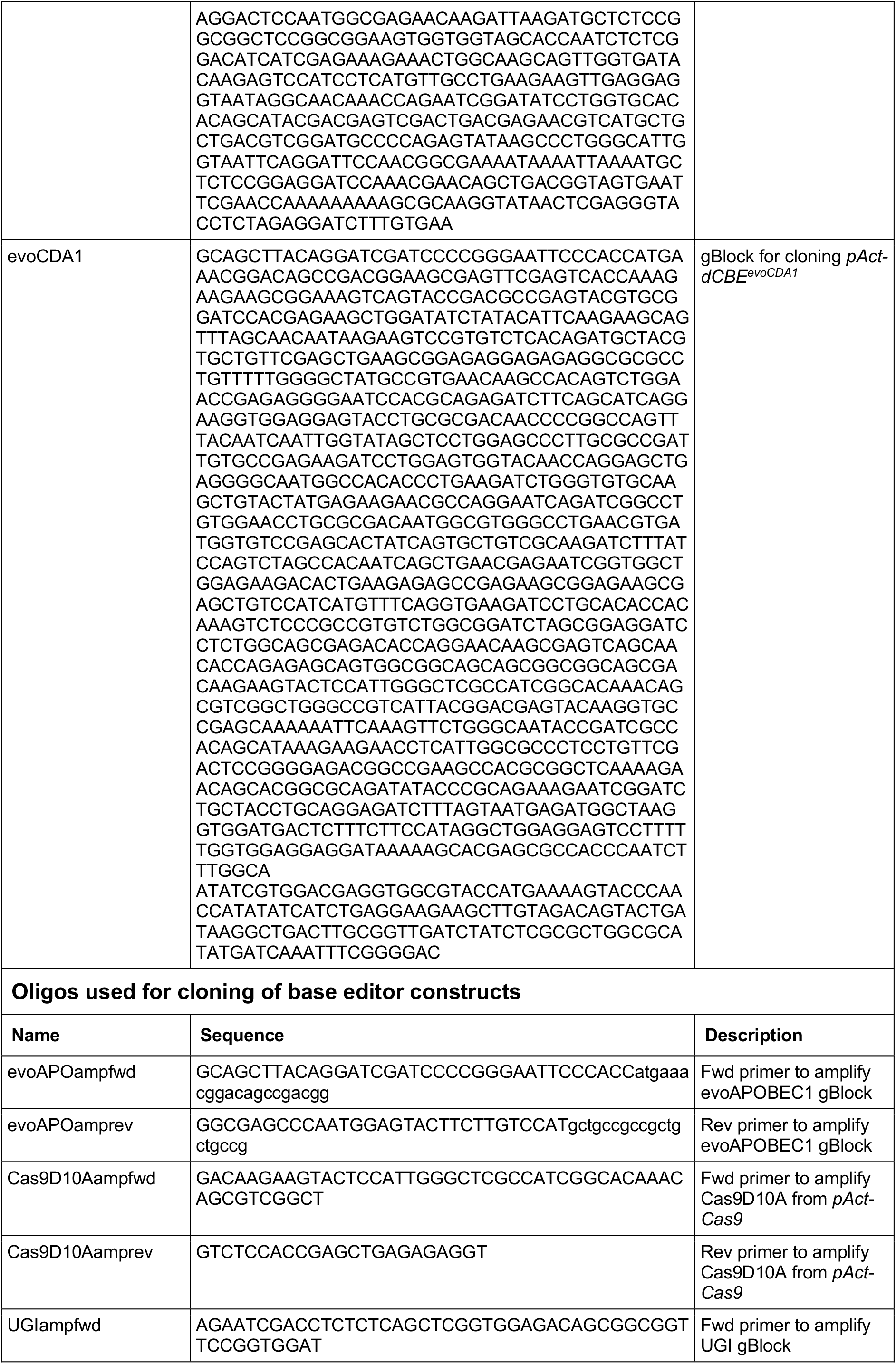
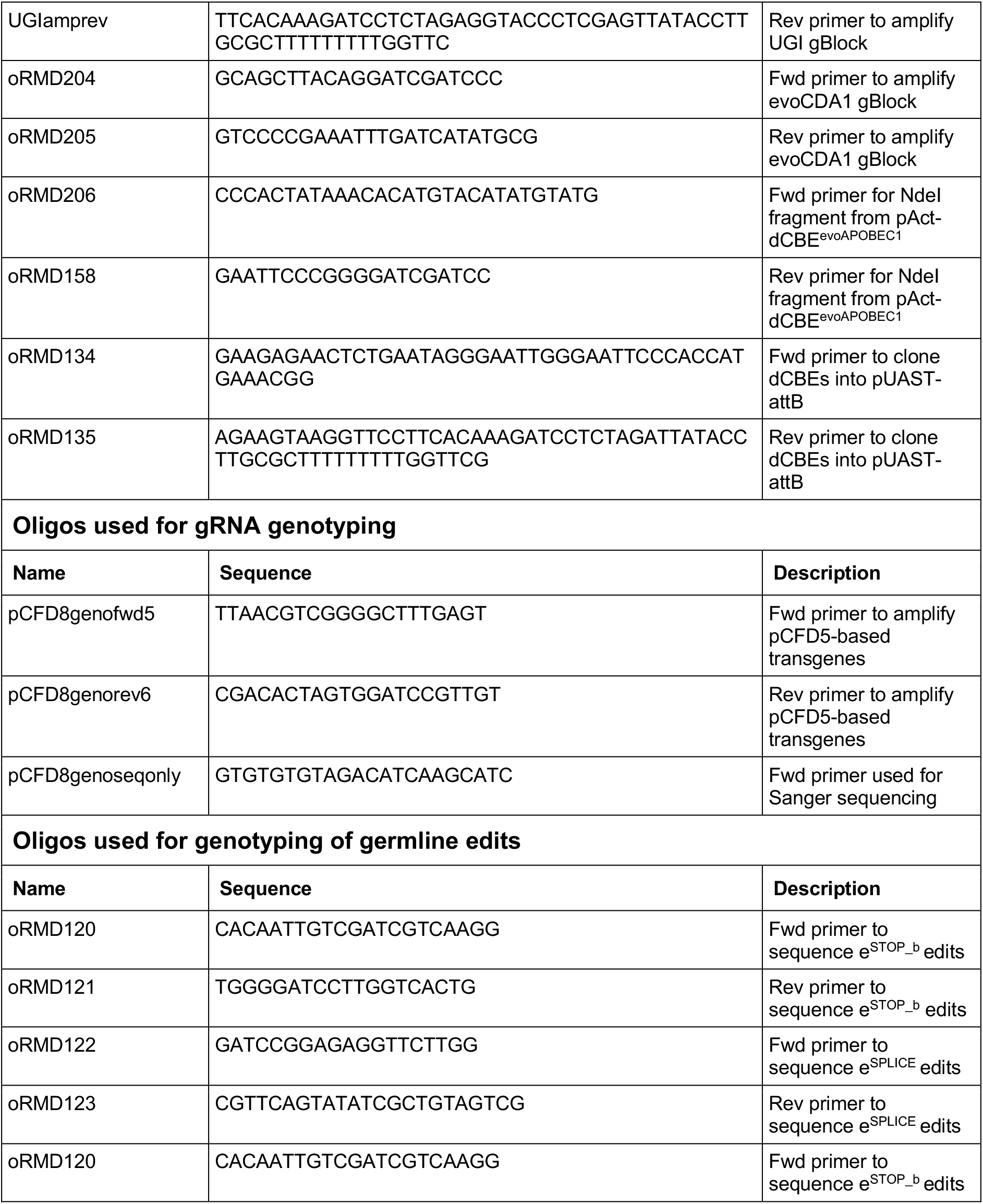
dsDNA fragments and oligonucleotides used in this work.

**Supplementary Table 2.**
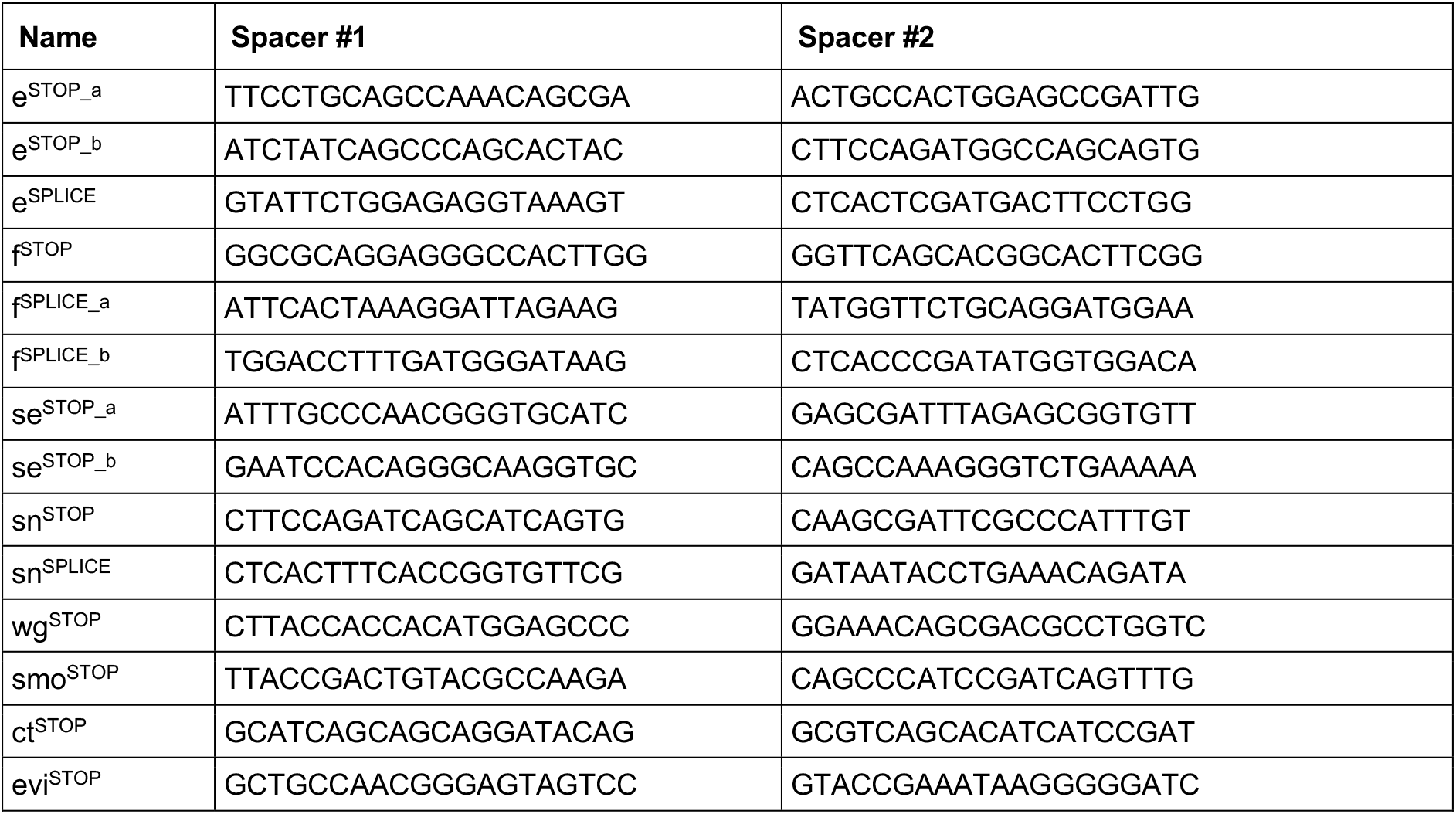
gRNA spacer sequences.

**Supplementary Table 3.**
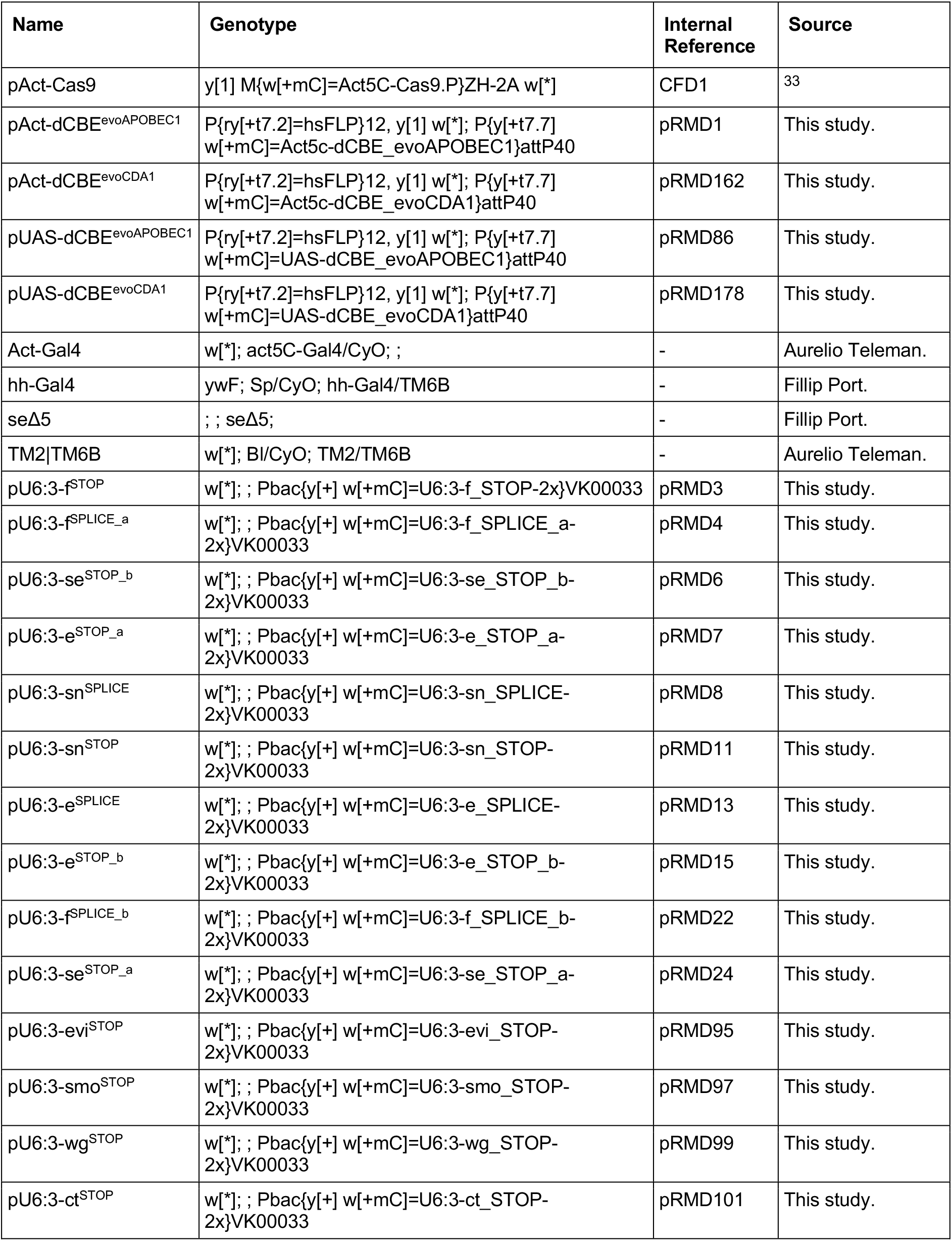
Fly strains used in this work.

| Name | Genotype | Internal Reference | Source |
| --- | --- | --- | --- |
| pAct-Cas9 | y[1] M{w[+mC]=Act5C-Cas9.P}ZH-2A w[*] | CFD1 | 33 |
| pAct-dCBE<sup>evoAPOBEC1</sup> | P{ry[+t7.2]=hsFLP}12, y[1] w[*]; P{y[+t7.7] w[+mC]=Act5c-dCBE_evoAPOBEC1}attP40 | pRMD1 | This study. |
| pAct-dCBE<sup>evoCDA1</sup> | P{ry[+t7.2]=hsFLP}12, y[1] w[*]; P{y[+t7.7] w[+mC]=Act5c-dCBE_evoCDA1}attP40 | pRMD162 | This study. |
| pUAS-dCBE<sup>evoAPOBEC1</sup> | P{ry[+t7.2]=hsFLP}12, y[1] w[*]; P{y[+t7.7] w[+mC]=UAS-dCBE_evoAPOBEC1}attP40 | pRMD86 | This study. |
| pUAS-dCBE<sup>evoCDA1</sup> | P{ry[+t7.2]=hsFLP}12, y[1] w[*]; P{y[+t7.7] w[+mC]=UAS-dCBE_evoCDA1}attP40 | pRMD178 | This study. |
| Act-Gal4 | w[*]; act5C-Gal4/CyO; ; | - | Aurelio Teleman. |
| hh-Gal4 | ywF; Sp/CyO; hh-Gal4/TM6B | - | Fillip Port. |
| seΔ5 | ; ; seΔ5; | - | Fillip Port. |
| TM2\|TM6B | w[*]; Bl/CyO; TM2/TM6B | - | Aurelio Teleman. |
| pU6:3-f<sup>STOP</sup> | w[*]; ; Pbac{y[+] w[+mC]=U6:3-f_STOP-2x}VK00033 | pRMD3 | This study. |
| pU6:3-f<sup>SPLICE_a</sup> | w[*]; ; Pbac{y[+] w[+mC]=U6:3-f_SPLICE_a-2x}VK00033 | pRMD4 | This study. |
| pU6:3-se<sup>STOP_b</sup> | w[*]; ; Pbac{y[+] w[+mC]=U6:3-se_STOP_b-2x}VK00033 | pRMD6 | This study. |
| pU6:3-e<sup>STOP_a</sup> | w[*]; ; Pbac{y[+] w[+mC]=U6:3-e_STOP_a-2x}VK00033 | pRMD7 | This study. |
| pU6:3-sn<sup>SPLICE</sup> | w[*]; ; Pbac{y[+] w[+mC]=U6:3-sn_SPLICE-2x}VK00033 | pRMD8 | This study. |
| pU6:3-sn<sup>STOP</sup> | w[*]; ; Pbac{y[+] w[+mC]=U6:3-sn_STOP-2x}VK00033 | pRMD11 | This study. |
| pU6:3-e<sup>SPLICE</sup> | w[*]; ; Pbac{y[+] w[+mC]=U6:3-e_SPLICE-2x}VK00033 | pRMD13 | This study. |
| pU6:3-e<sup>STOP_b</sup> | w[*]; ; Pbac{y[+] w[+mC]=U6:3-e_STOP_b-2x}VK00033 | pRMD15 | This study. |
| pU6:3-f<sup>SPLICE_b</sup> | w[*]; ; Pbac{y[+] w[+mC]=U6:3-f_SPLICE_b-2x}VK00033 | pRMD22 | This study. |
| pU6:3-se<sup>STOP_a</sup> | w[*]; ; Pbac{y[+] w[+mC]=U6:3-se_STOP_a-2x}VK00033 | pRMD24 | This study. |
| pU6:3-evi<sup>STOP</sup> | w[*]; ; Pbac{y[+] w[+mC]=U6:3-evi_STOP-2x}VK00033 | pRMD95 | This study. |
| pU6:3-smo<sup>STOP</sup> | w[*]; ; Pbac{y[+] w[+mC]=U6:3-smo_STOP-2x}VK00033 | pRMD97 | This study. |
| pU6:3-wg<sup>STOP</sup> | w[*]; ; Pbac{y[+] w[+mC]=U6:3-wg_STOP-2x}VK00033 | pRMD99 | This study. |
| pU6:3-ct<sup>STOP</sup> | w[*]; ; Pbac{y[+] w[+mC]=U6:3-ct_STOP-2x}VK00033 | pRMD101 | This study. |

**Supplementary Table 4.**
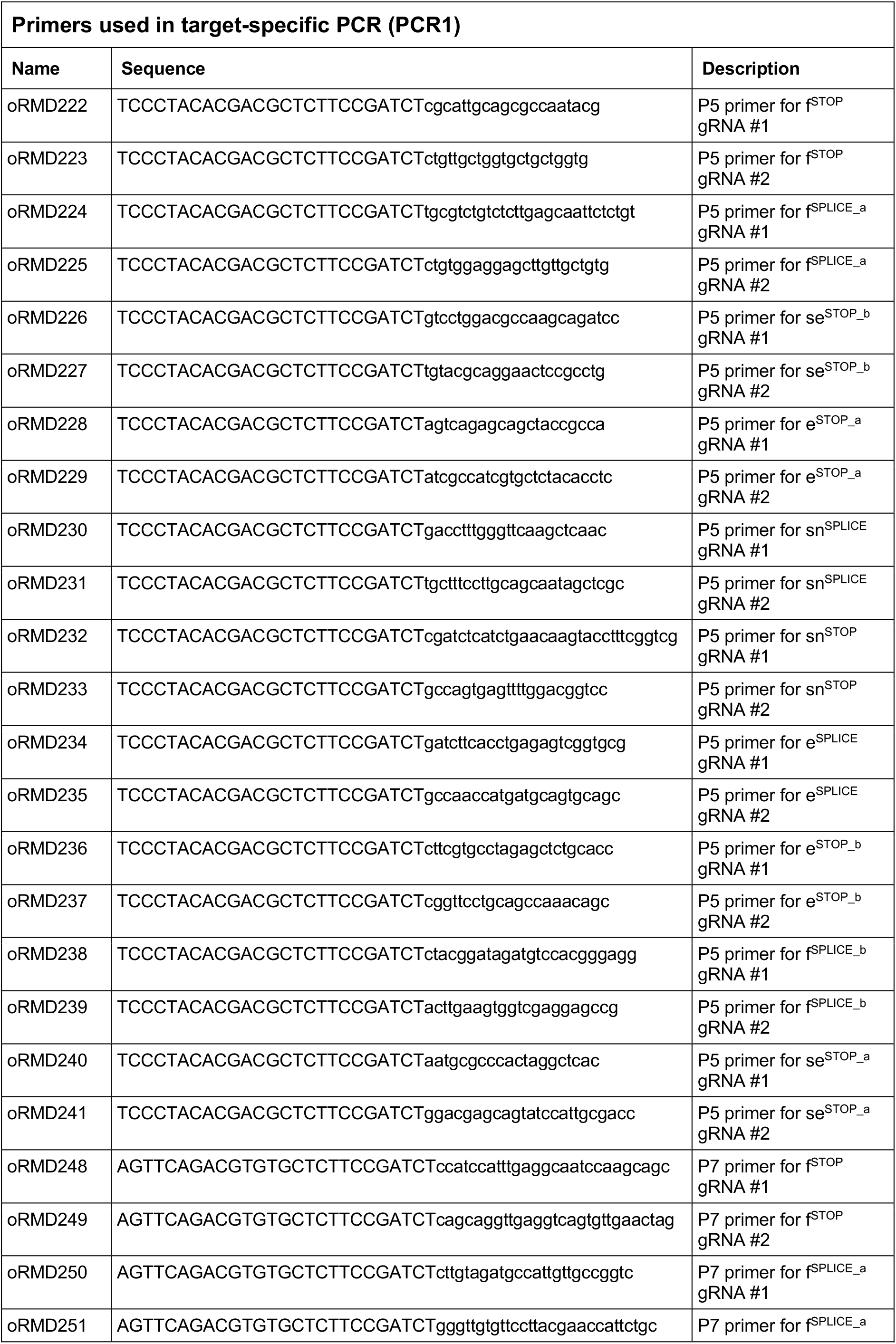
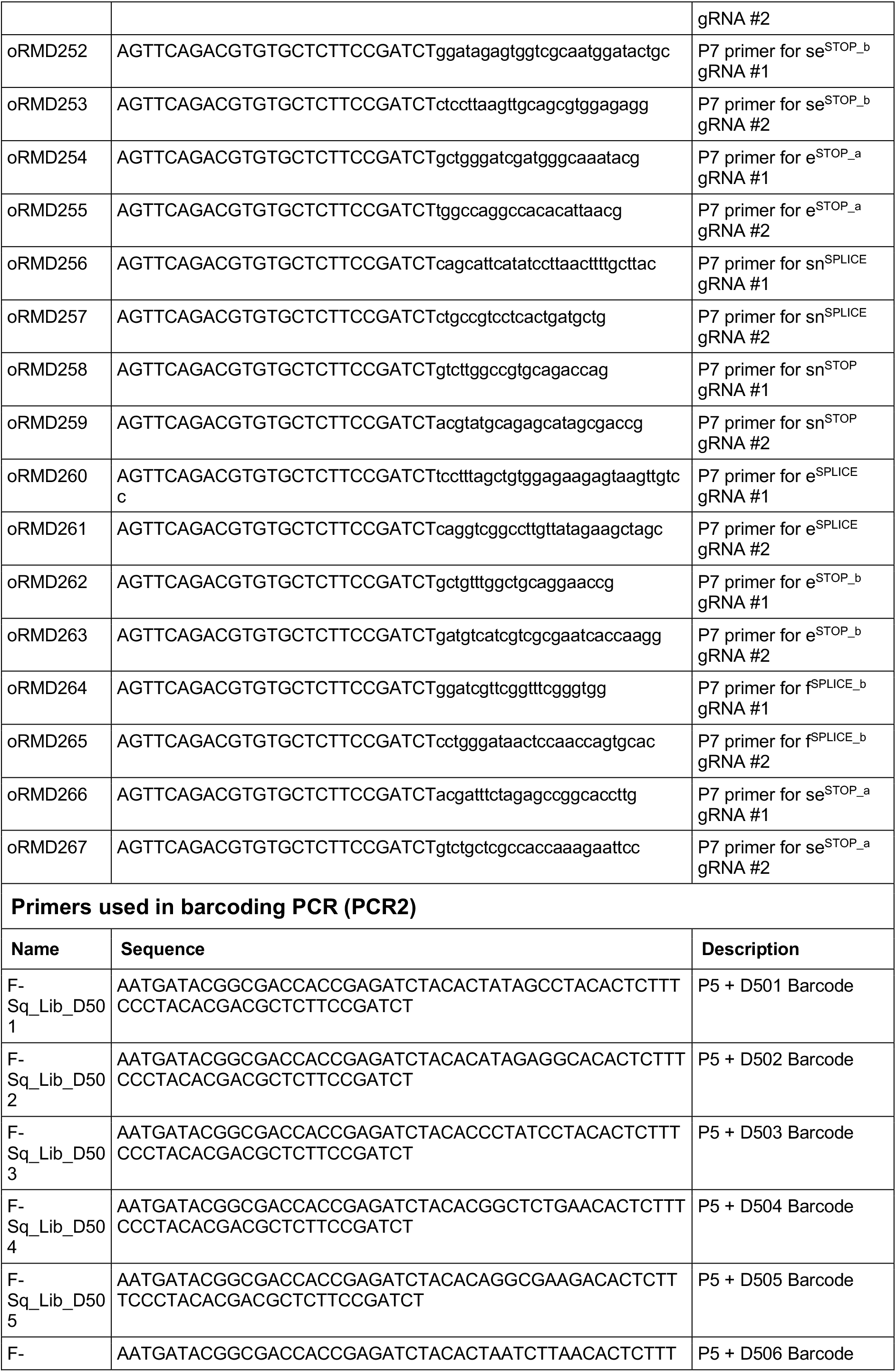
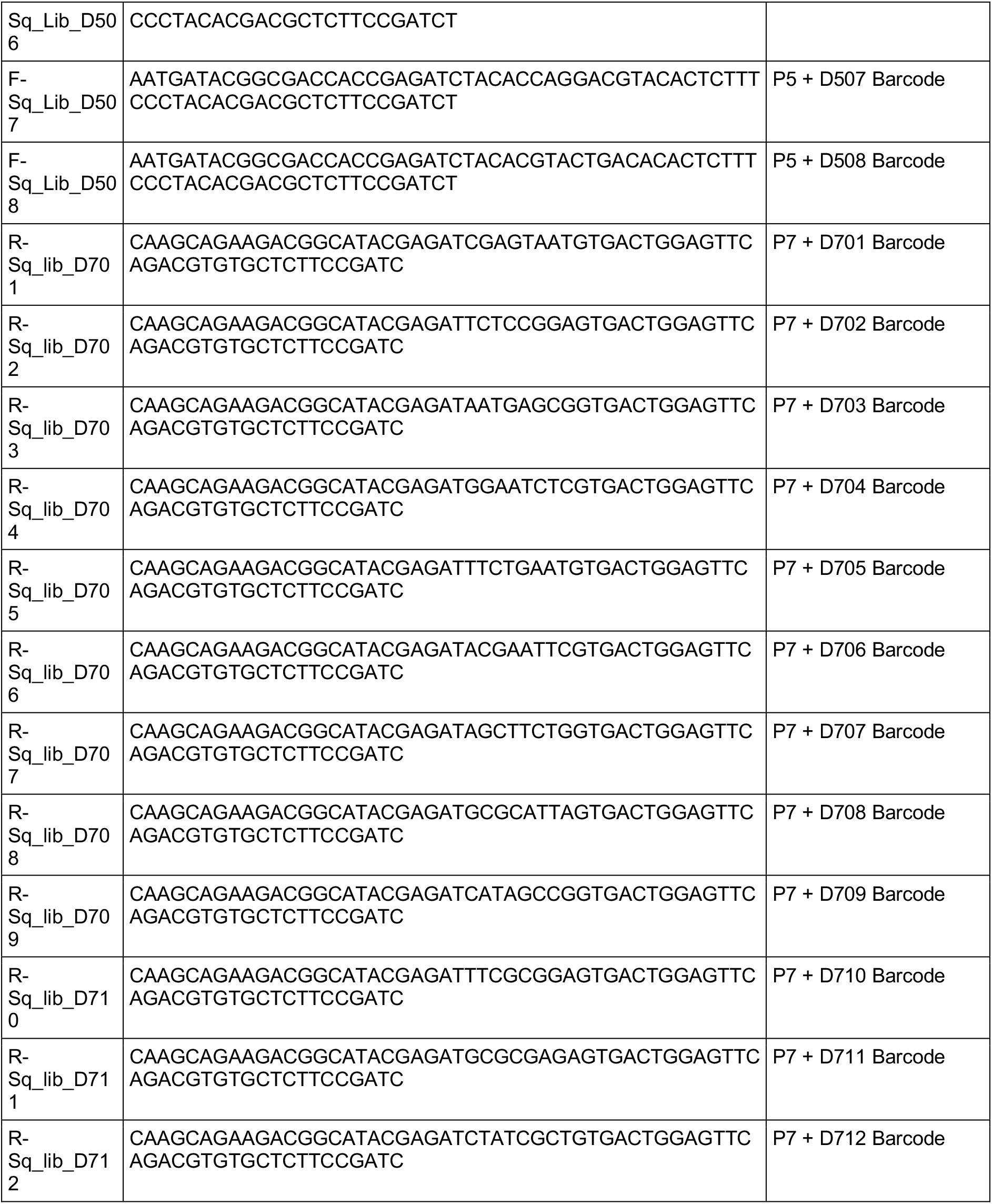
Primers used for Amp-Seq library preparation.

**Supplementary Table 5.**
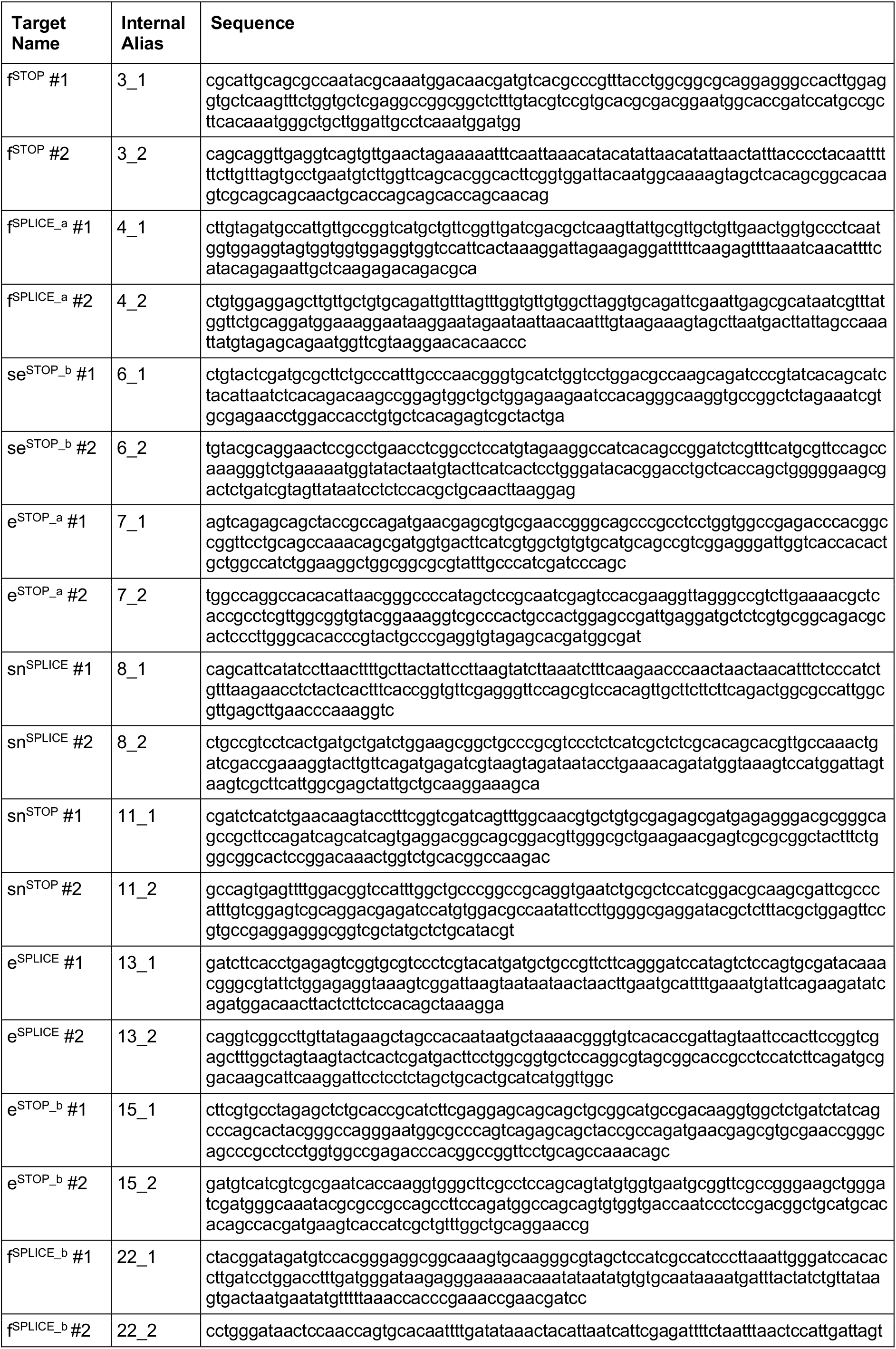
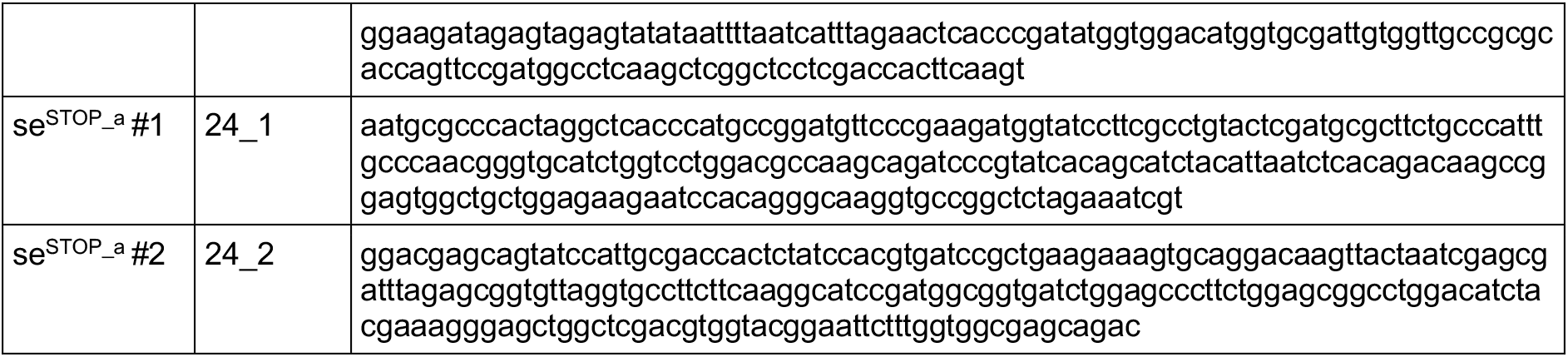
Sequences of genomic amplicons subjected to Amp-Seq.

## Supplementary Figures

**Supplementary Figure 1.**
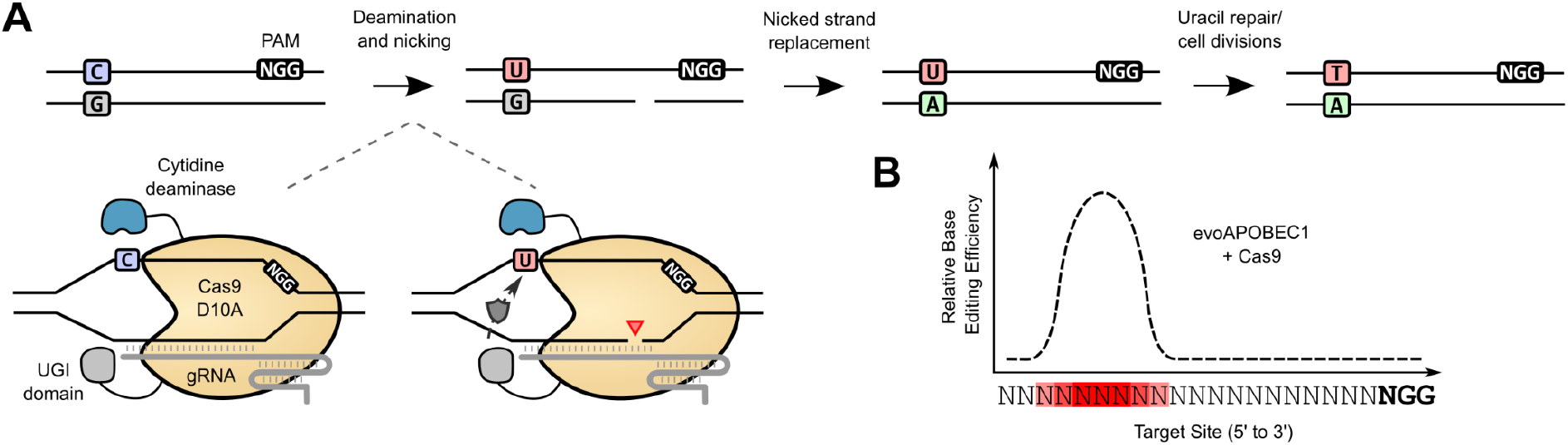
Schematic representations of the cytosine base editing mechanism. **(A)** Following deamination of cytosine residues and nicking of the target strand, endogenous DNA repair pathways and cell divisions establish the C:G to T:A edit. **(B)** Canonical editing window of CBEs utilising Cas9 and the evoAPOBEC1 domain. Efficient C-to-T editing is generally observed at positions 3-9 of the protospacer, counting from 5’.

**Supplementary Figure 2.**
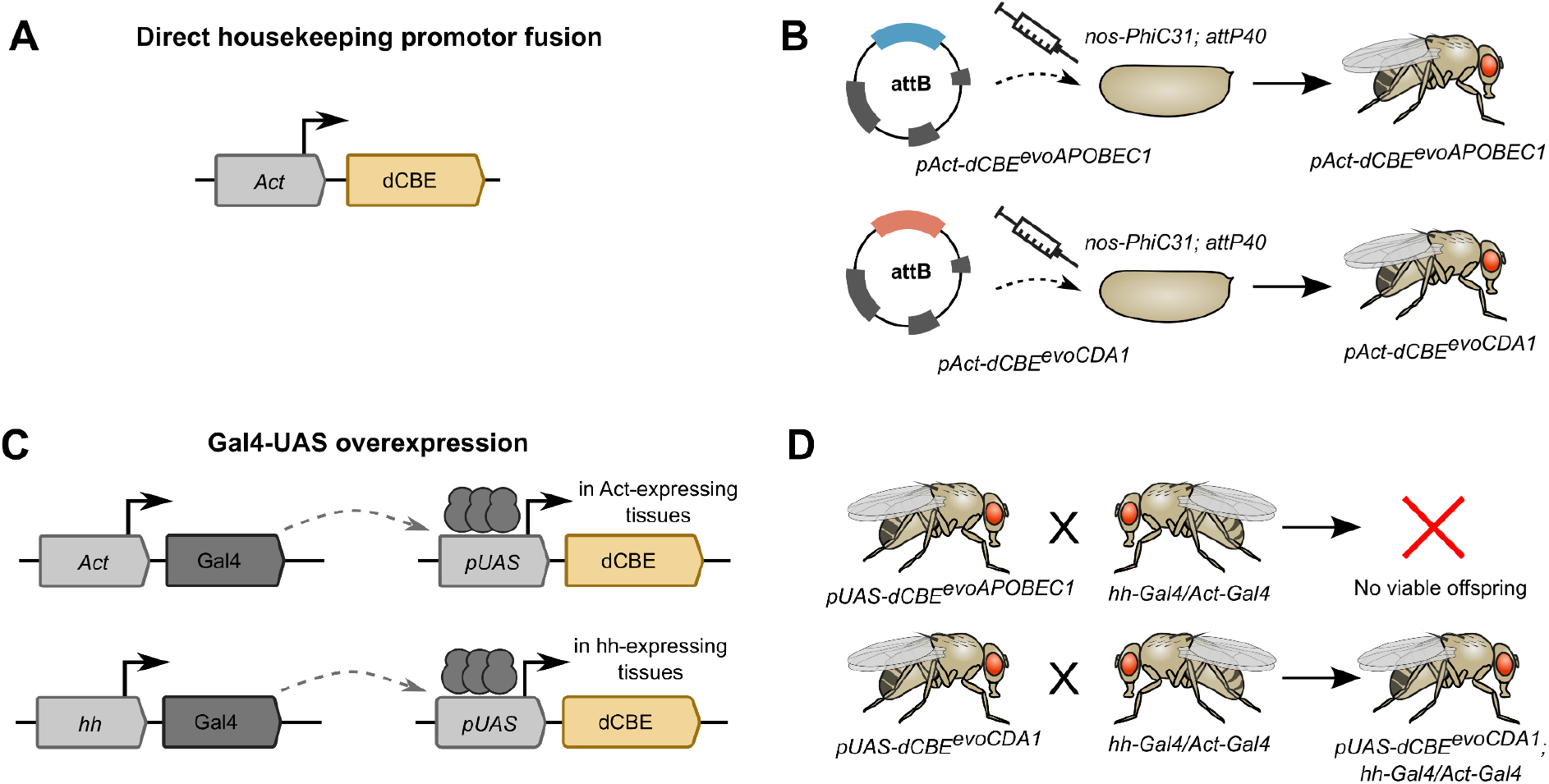
Generation of dCBE-expressing transgenic *Drosophila* strains. **(A)** Schematic representation of a direct promoter fusion. The base editor is placed directly downstream of the promoter of the housekeeping gene *act5c* (*pAct*). **(B)** Generation of stable transgenic stocks for *pAct-dCBE^evoAPOBEC1^* and *pAct-dCBE^evoCDA1^*. For both editors, viable and fertile stocks could be obtained following embryo microinjection. **(C)** Schematic representation of the Gal4-UAS expression system. A regulatory sequence of choice drives Gal4 expression, which then activates transcription from UAS promoters in the respective tissues. **(D)** Induction of base editor expression by crosses to Gal4 driver lines. Viable offspring in crosses with *hh-Gal4* and *Act-Gal4* driver lines was only obtained with *pUAS-dCBE^evoCDA1^*.

**Supplementary Figure 3.**
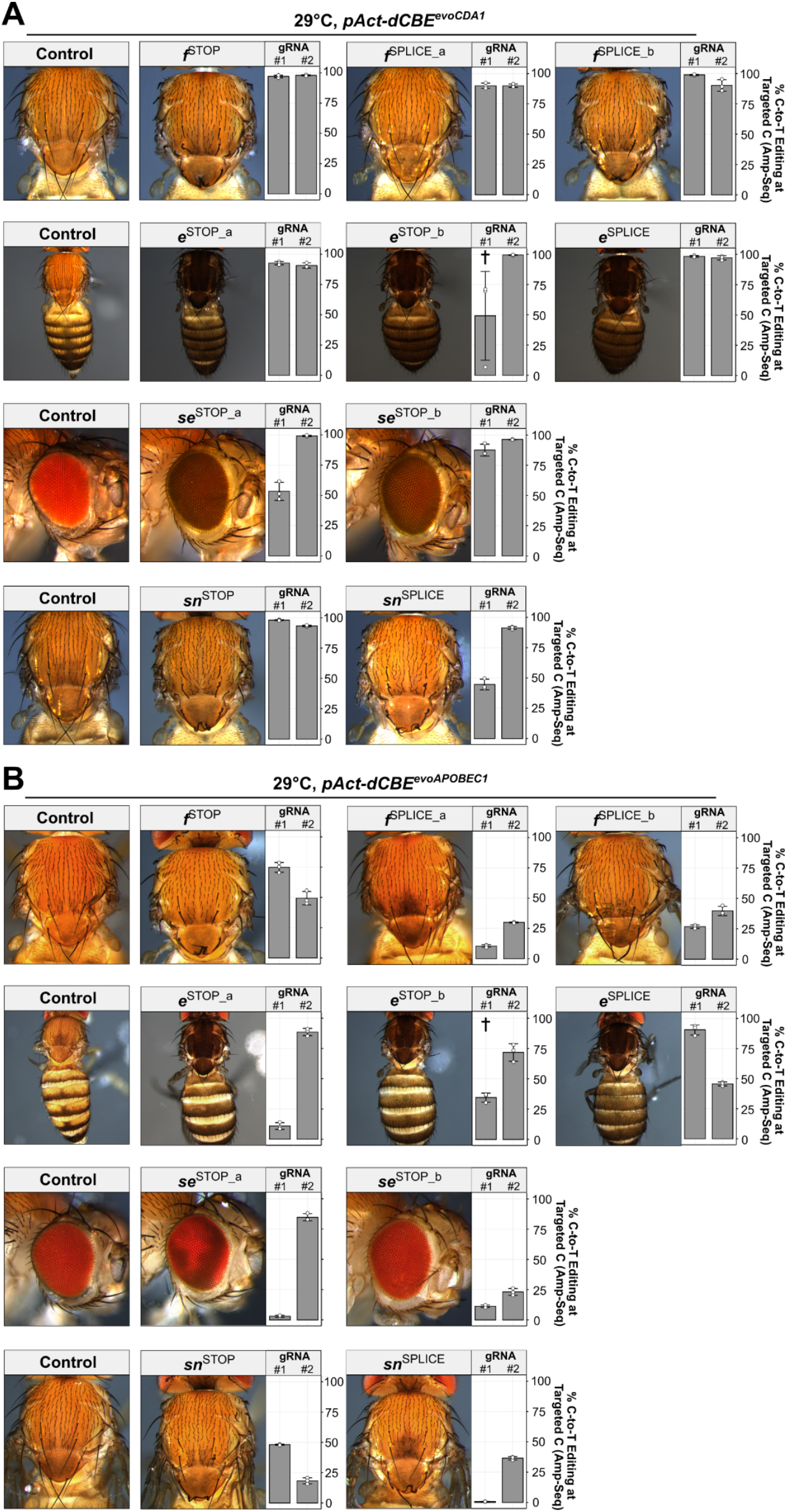
Mutagenesis of non-essential genes with dCBE^evoCDA1^ (A) and dCBE^evoAPOBEC1^ (B) at 29°C. Representative images of female offspring are shown alongside the C-to-T editing rates at the targeted C residues, determined by Amp-Seq (n = 3, data presented as mean ± s.d. (bars and error bars) and individual measurements (points)). For all strains except *f*^SPLICE_a^, fully penetrant loss-of-function phenotypes are observed with dCBE^evoCDA1^, whereas mutagenesis efficiency is highly variable with dCBE^evoAPOBEC1^. The C-to-T editing rates reflect the severity of the observed phenotype.

**Supplementary Figure 4.**
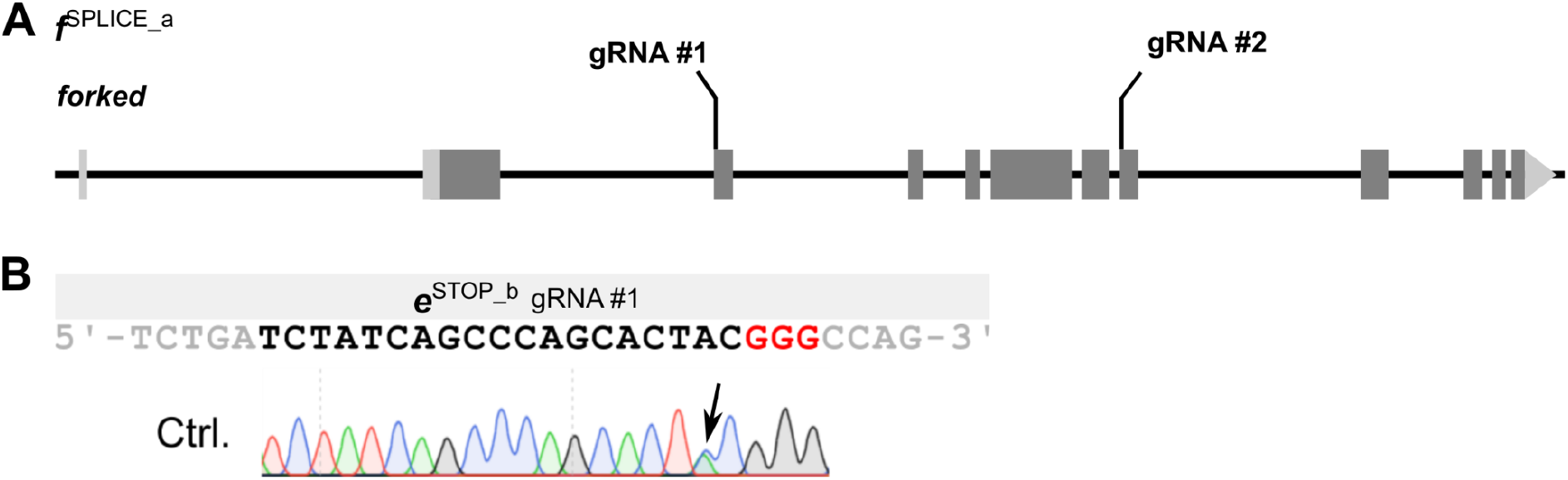
Additional information on target sites. **(A)** Schematic representation of the *forked* gene and the positions of the target sites of *f*^SPLICE_a^. Both gRNAs target the splice acceptors of small exons. **(C)** Sanger sequencing of target site of the *e*^STOP_b^ gRNA #1. A A/C SNP is detected two nucleotides upstream of the PAM (in red).

**Supplementary Figure 5.**
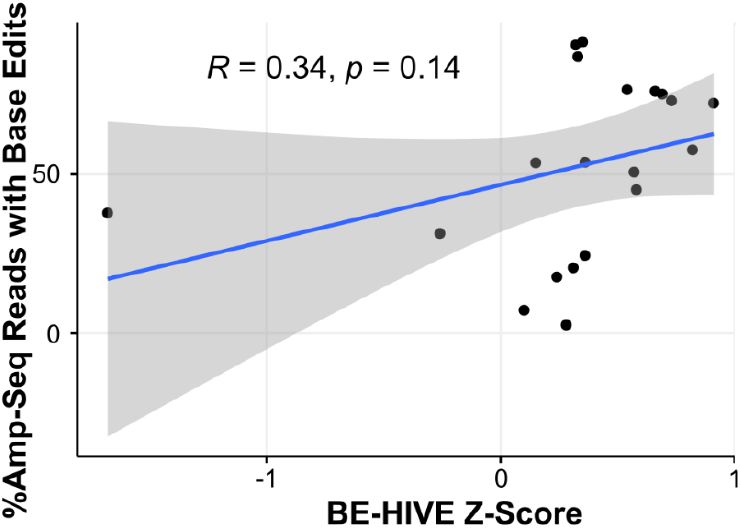
Correlation of dCBE^evoAPOBEC1^ activity with BE-HIVE prediction. Each dot represents one gRNA (N = 20). A modest but non statistically significant correlation is observed (Pearson correlation).

**Supplementary Figure 6.**
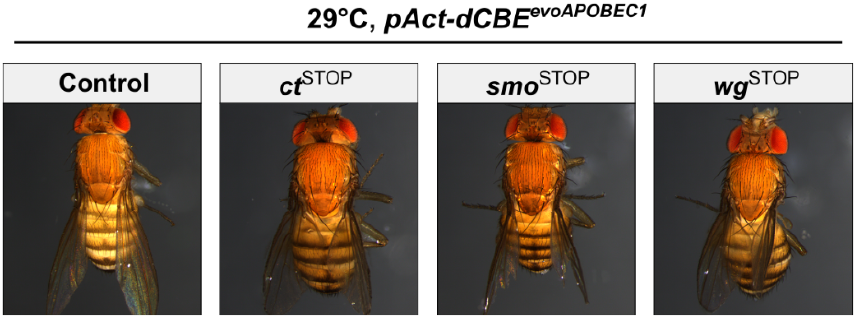
Mutagenesis of the essential target genes with dCBE^evoAPOBEC1^ at 29°C. Representative images of female offspring from crosses with *pAct-dCBE^evoAPOBEC1^* against gRNAs targeting the essential genes *ct*, *smo* and *wg* raised at 29°C are shown. In contrast to *pAct-dCBE^evoCDA1^*, the crosses result in viable offspring which exhibit minor wing malformations.

**Supplementary Figure 7.**
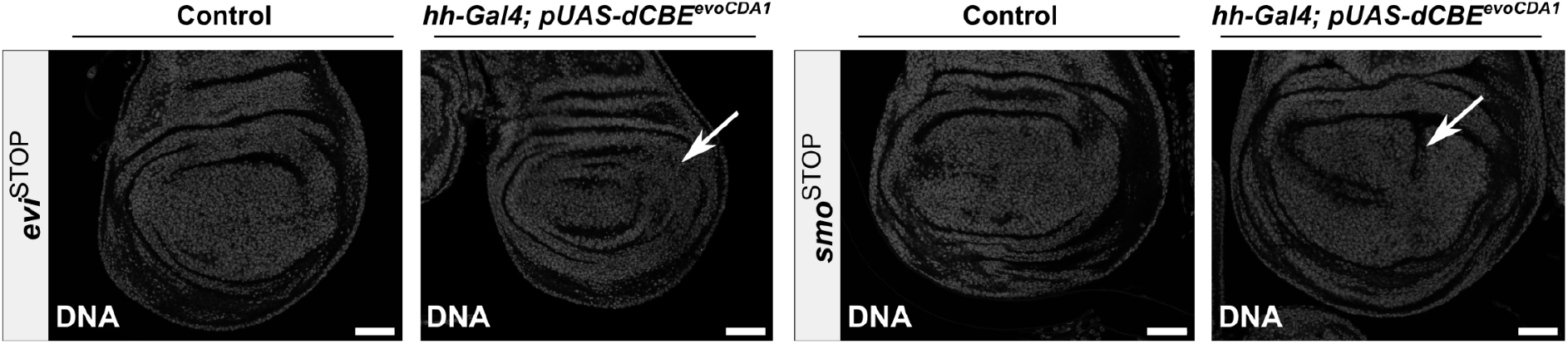
DNA staining of wing imaginal discs subjected to mutagenesis with *hh-Gal4*; *pUAS-dCBE^evoCDA1^*. Compared to control conditions, instances of morphological abnormalities are detected in the base editor expressing compartment (white arrows). Scale bars represent 50 μm.

**Supplementary Figure 8.**
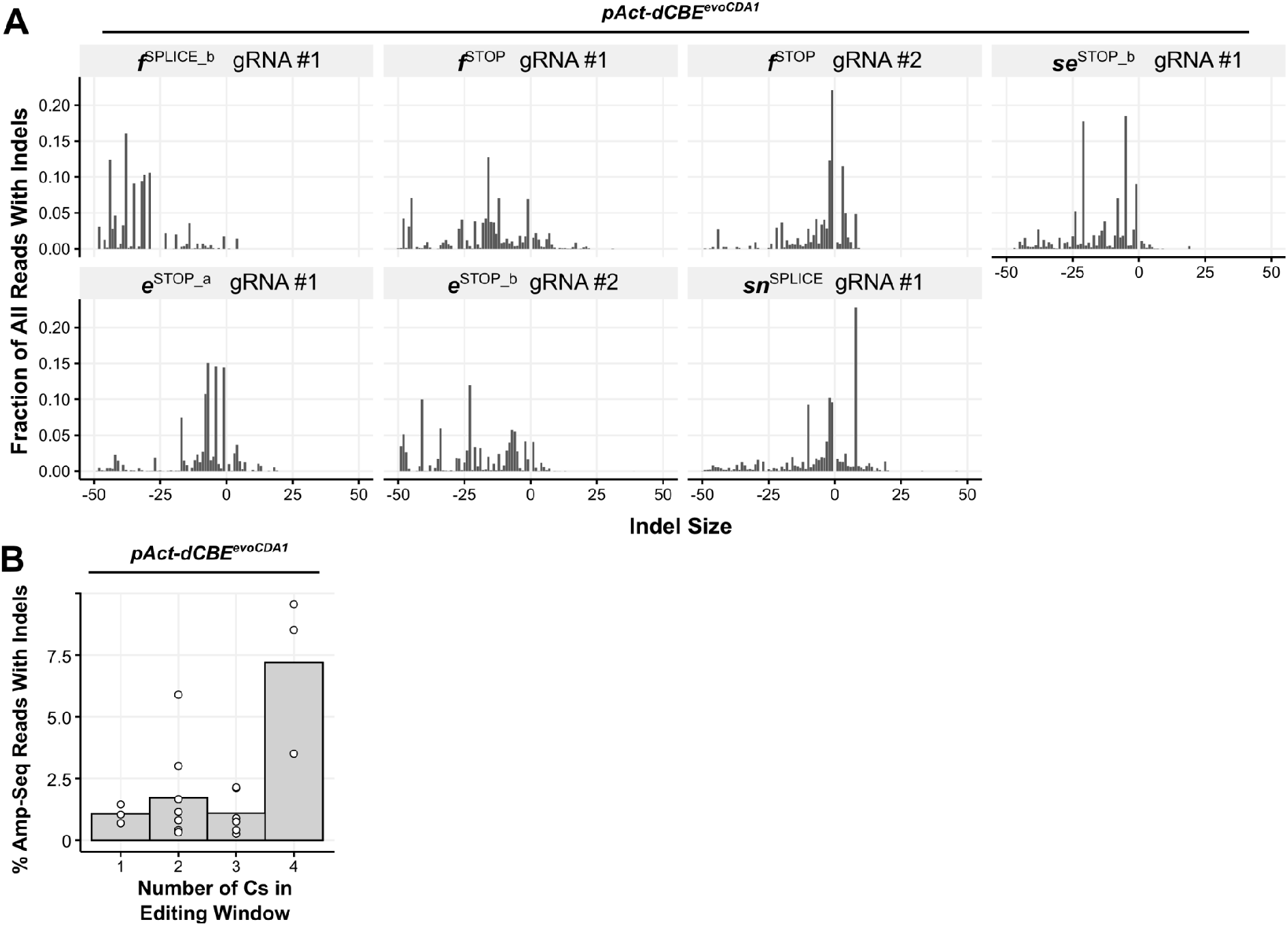
Analysis of indel by-products. **(A)** Indel size distribution with dCBE^evoCDA1^ at sites exhibiting more than 2% Amp-Seq reads with indels (data presented as fraction of all reads containing indels, averaged across n = 3 replicates). A broad range of insertion and deletion sizes is observed. **(B)** % Amp-Seq reads containing indels as a function of the number of C residues in the editing window (data presented as mean (bars) and individual measurements (each dot representing the average indel percentage from n = 3 replicates for one target site). Sites with four C residues in their editing window tend to exhibit higher indel formation rates.

